# A hyperspherical deep Bayesian model for interpretable clustering and relationship prediction in microbiome multi-omics integration

**DOI:** 10.64898/2026.07.28.741374

**Authors:** Tung Dang, Artem Lysenko, Tatsuhiko Tsunoda

## Abstract

The microbiome plays a significant role in the development and progression of many diseases, yet extracting interpretable insights from multi-omics data remains challenging. Existing approaches face a recurring practical trade-off: deep learning methods achieve high predictive performance but lack uncertainty quantification, whereas probabilistic methods provide interpretable results but require data-type-specific likelihood functions that limit generalization across diverse omics modalities. Here, we introduce DBayesCM (Deep Bayesian Clustering for Multi-omics), which combines deep learning modeling with Bayesian nonparametric methods. DBayesCM employs separate encoders to project microbiome and host omics data into a shared latent space, where an infinite mixture model with a Dirichlet process prior determines the number of clusters automatically while quantifying the uncertainty of each sample’s assignment. Spike-and-slab priors identify discriminative features, and a Bayesian neural network estimates probabilistic co-occurrence between microbial species and host omics features. To isolate the effect of latent geometry, we evaluate two variants that are identical except for their latent space: DBayesCM-vMF constrains the latent to the unit hypersphere and applies a von Mises-Fisher mixture, while DBayesCM-GMM uses a Euclidean latent space and a Gaussian mixture. On simulated data, the hyperspherical variant recovered the correct number of clusters, whereas the Euclidean variant over-segmented, demonstrating that the latent geometry affects cluster recovery. Applied to colon, breast, and kidney cancer cohorts spanning metagenomics, host metabolomics, RNA-seq, and miRNA data, and to an obstructive sleep apnea model, DBayesCM ranked consistently among the existing methods while uniquely combining data-driven cluster-number determination, sample-level uncertainty, and interpretable feature selection within a single framework. DBayesCM reveals conditional probabilistic co-occurrence between core microbial species and host omics features, enabling uncertainty-aware exploration of microbiome-host relationships across diverse diseases.

## Introduction

The human microbiome profoundly influences health and disease through complex interactions with host metabolism, immunity, and cellular signaling [1, 2]. Integrating microbiome profiling (metagenomics) with host multiomics data (metabolomics, RNA-seq, and miRNA) promises to reveal disease mechanisms and enable precision patient stratification [3, 4]. Such integration requires methods that can discover clusters from heterogeneous data modalities, quantify prediction uncertainty for clinical decision-making, and identify interpretable biomarkers with mechanistic insights [5, 6]. However, existing methods face an impactful practical trade-off between clustering accuracy and probabilistic interpretability.

Deep learning approaches for multiomics integration, such as SnapCCESS [7] and MultiVI [8] employ variational autoencoders to achieve superior clusters discovery but provide only point estimates without uncertainty quantification. This is a critical limitation when clinical decisions depend on distinguishing reliable classifications from ambiguous cases that require additional investigation [9]. Dimensionality reduction methods, such as MOFA+ [10] use variational inference to learn latent factors from multiomics data but extract expectations of variational distributions for downstream analysis rather than providing full posterior distributions over cluster assignments. Moreover, users must specify the number of factors in advance and perform separate clustering on learned factors. Integrative clustering methods, such as iCluster [11] jointly model modalities through latent variables with generalized linear models (Poisson distribution for counts), but select the number of clusters through a separate model-selection procedure and computationally expensive gene preselection via EM-based lasso regularization [12], rather than inferring cluster number and its uncertainty within a single model. Specialized microbiome clustering methods, such as SVVS [13] provide automatic cluster discovery and uncertainty quantification through an infinite Dirichlet multinomial mixture model. However, it requires separate mixture models for each data type, limiting its direct extension to heterogeneous multiomics integration. Bayesian integrative methods such as iClusterBayes [14] and Bayesian consensus clustering [15] provide posterior uncertainty over cluster assignments, but do not incorporate the deep encoders needed to model the non-linear, complex structure of integrated microbiome-omics data, nor sample-level uncertainty coupled with automatic cluster-number discovery.

Here, we present DBayesCM (Deep Bayesian Clustering for Multi-omics), a Bayesian deep learning framework that integrates microbiome and host multi-omics data to achieve uncertainty-aware clustering with automatic cluster-number discovery and disease-specific cross-modality dependency mapping. DBayesCM addresses four fundamental challenges simultaneously: (1) learning unified representations from heterogeneous count modalities (microbiome, metabolomics, RNA-seq, and miRNA); (2) clustering samples while inferring the number of clusters from the data; (3) quantifying uncertainty in both sample assignments and feature selection; and (4) uncovering probabilistic relationships between microbial species and host molecular profiles.

In this study, all integrated modalities are represented as count data. DBayesCM therefore employs modality-specific variational autoencoders with Dirichlet-multinomial reconstruction likelihoods for every modality, explicitly modeling the overdispersion and compositionality inherent in count-based omics data. These learned representations are fused through a shared latent space for joint analysis, which an infinite mixture with a Dirichlet process prior then clusters, discovering the number of clusters from the data while providing full posterior distributions over sample assignments. To isolate the effect of the latent geometry on clustering, we evaluate two variants that are identical in architecture, prior structure, and training, and differ only in how the shared latent space is modeled. DBayesCM-vMF constrains the latent to the unit hypersphere and clusters it with a von Mises-Fisher mixture, so that samples are grouped by the relative shape of their multi-omic profile rather than by overall magnitude, a property suited to compositional data, where relative composition is more informative than absolute abundance [16, 17, 18]. DBayesCM-GMM uses a Euclidean latent and a Gaussian mixture [19]. Because the two variants share all other components, their comparison directly quantifies the contribution of the hyperspherical geometry, a controlled comparison that is not possible against external methods, which differ from DBayesCM in encoder, likelihood, and inference simultaneously. For feature selection, DBayesCM employs spike-and-slab shrinkage priors [20] that identify a minimal core set of features across all modalities while quantifying the selection confidence through posterior probabilities. Additionally, DBayesCM integrates cluster-specific Bayesian neural networks (VBayesMM) [21] to estimate cluster-specific conditional associations between microbial species and host omics features, providing a probabilistic interpretation of how microbiome and host molecular profiles co-vary across clusters.

We evaluated DBayesCM on multiomics microbiome datasets from published case-control cohorts (colorectal cancer [22] and obstructive sleep apnea [23]) and integrated TCGA-TCMA databases (kidney and breast cancers [24, 25]). DBayesCM consistently outperformed SnapCCESS, MultiVI, MOFA+, and iCluster in terms of clustering accuracy, while automatically identifying correct cluster numbers and providing sample-level uncertainty quantification.

## Materials and methods

### The DBayesCM approach

#### Overview

DBayesCM combines deep learning modeling with Bayesian nonparametric feature selection, thus inheriting the benefits of flexible nonlinear modeling and interpretability. It considers cancer microbiome abundances and host multiomics data (metabolite, RNA-seq, and miRNA) and outputs probabilistic cluster assignments with quantified uncertainty, visualization of the recovered cluster geometry, feature selection with posterior inclusion probabilities, and co-occurrence relationships between microbiome and omics features (Fig. 1).

**Fig. 1.**
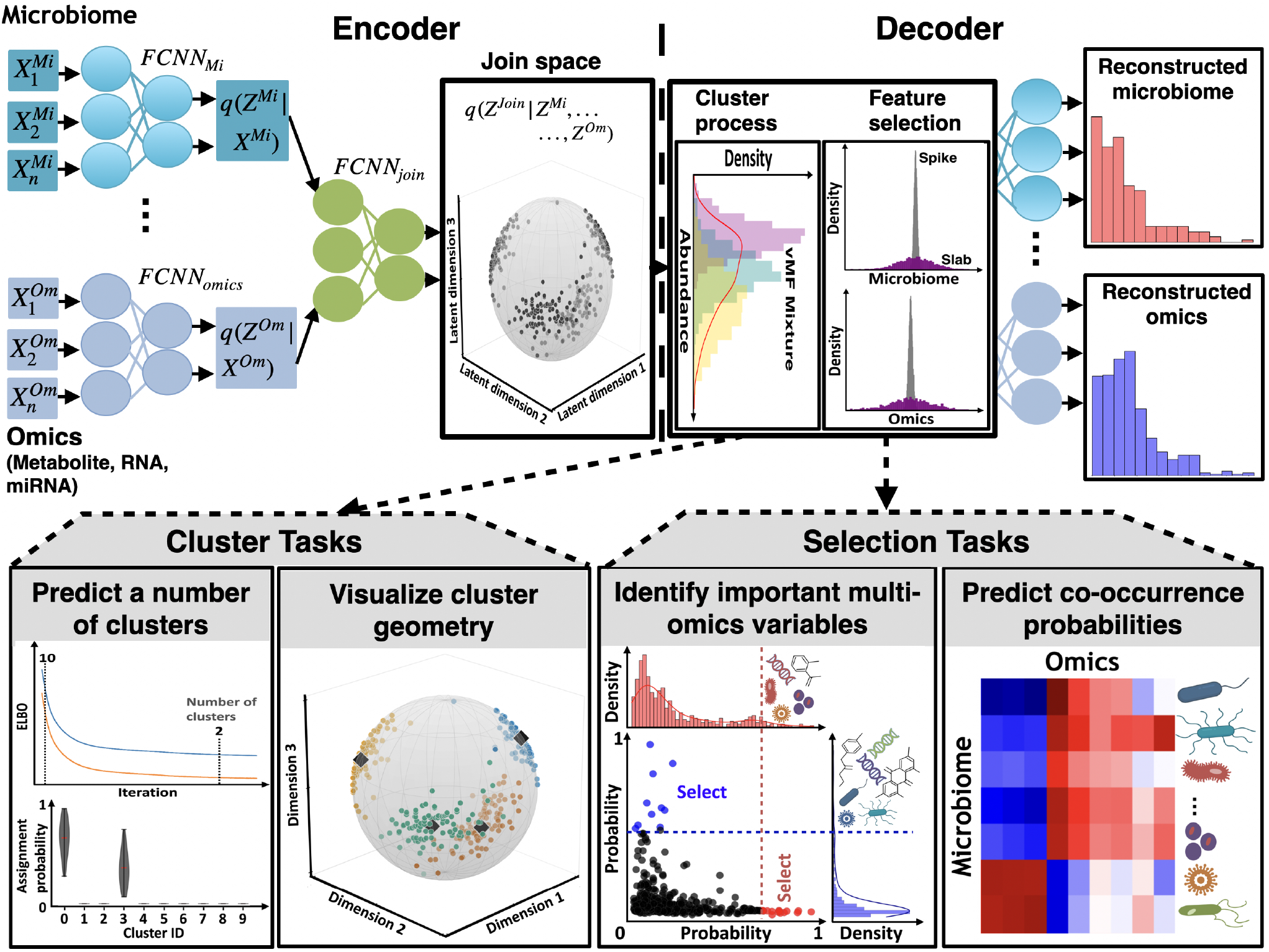
Schematic of the DBayesCM model (DBayesCM-vMF shown). DBayesCM integrates microbiome (*X*^*Mi*^) and host omics (*X*^*Om*^) data (metabolite, RNA, miRNA) through a deep Bayesian architecture. **Encoder**: modality-specific fully connected neural networks (FCNNs) generate latent representations *Z*^*Mi*^ and *Z*^*Om*^, fused into a joint latent space *Z*^*Join*^ through a fusion layer (FCNN_*join*_), constrained to the unit hypersphere in DBayesCM-vMF (DBayesCM-GMM uses a Euclidean latent). **Decoder**: (1) an infinite mixture model with a Dirichlet process prior, parameterized as a von Mises-Fisher mixture (or a Gaussian mixture of DBayesCM-GMM), assigns samples to clusters and infers the number of clusters from the data while quantifying assignment uncertainty, (2) spike-and-slab shrinkage priors select discriminative microbiome and omics features. The model reconstructs microbiome and omics data via Dirichlet-multinomial distributions. The DBayesCM supports four downstream applications: (1) predicting the optimal number of clusters together with quantifying assignment uncertainty, (2) visualizing the recovered cluster geometry, (3) identifying important multiomics variables, and (4) predicting microbiome-omics co-occurrence probabilities via Bayesian neural network.

In its core architecture, DBayesCM models integration through three components. First, separate encoders (fully connected neural networks, FCNNs) process microbiome data (*X*^*Mi*^) and omics data (*X*^*Om*^) into modality-specific latent representations *Z*^*Mi*^ and *Z*^*Om*^. These representations are combined into a unified joint latent space *Z*^*Join*^ that captures the cross-modal dependencies. Second, the decoder implements an infinite mixture model with a Dirichlet process prior that assigns samples to clusters, infers the number of clusters from the data, and provides posterior distributions over assignments. To isolate the effect of latent geometry, we evaluate two variants that are identical except for this component, DBayesCM-vMF, using a von Mises-Fisher mixture on a hyperspherical latent [16, 17], and DBayesCM-GMM, using a Gaussian mixture in Euclidean space [19]. In parallel, spike-and-slab shrinkage priors perform probabilistic feature selection [20], so that each cluster is characterized by a limited set of discriminative features, and each modality is reconstructed from Dirichlet-multinomial distributions [13] parameterized by cluster-specific selection matrices and cluster proportions. Third, a Bayesian neural network component adapted from the VBayesMM method [21] predicts co-occurrence probabilities between selected microbial species and omics features. DBayesCM is trained using variational inference by optimizing the evidence lower bound (ELBO) [19].

#### Generative model

DBayesCM integrates count-based multiomics data, including microbiome metagenomics, metabolomics, RNA-seq, and miRNA-seq, through a Bayesian model that explicitly models the compositional and overdispersed nature of omics count data. For the *n*^*th*^ sample (*n* ∈ [1, …, *N* ]),let 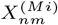 denote the read count for the *m*^*th*^ taxonomic unit (*m* ∈ [1, …, *M* ]) and 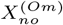 denote the count for the *o*^*th*^ host omics feature (*o* ∈ [1, …, *O*]), where *Om* ∈ *{*metabolomics, RNA-seq, miRNA-seq*}*. To capture compositional (features exist as proportions of total counts) and overdispersion (variance exceeds that of simple multinomial models), we model **X**^(*Mi*)^ and **X**^(*Om*)^ as Dirichlet-multinomial distributions parameterized by concentration parameters ***α*** and ***β*** respectively [13]:

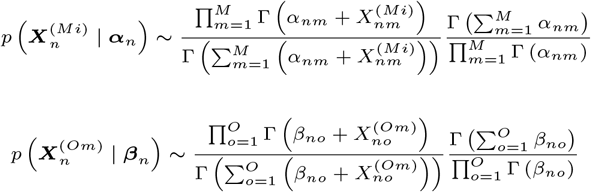

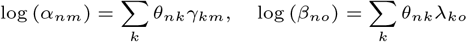

where Γ (.) is the gamma function. The key contribution of this study lies in decomposing the concentration parameters to enforce a shared latent clustering structure across modalities. Specifically, each sample parameter is expressed as a log-linear combination of cluster proportions *θ*_*nk*_ (the proportion of the *k*^*th*^ cluster in *n*^*th*^ sample) and cluster-specific feature embeddings:

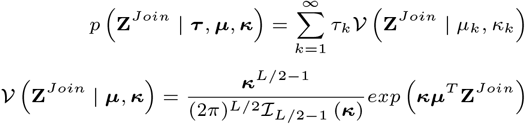

where *γ*_*km*_ represents the microbiome embedding of taxonomic unit *m*^*th*^ in *k*^*th*^ cluster and *λ*_*ko*_ represents the host omics embedding of feature *o*^*th*^ in *k*^*th*^ cluster. This approach ensures that samples sharing similar cluster memberships exhibit correlated microbiome and omics profiles, enabling the joint discovery of clusters across data modalities.

#### Encoders, joint latent space, and the two variants

A joint encoder produces shared latent representations 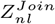 for sample *n* in latent dimension *l* (*l* ∈ [1, …, *L*]), encoding information across microbiome and host omics modalities. Modality-specific encoder neural networks (three fully connected layers with ReLU activations and dropout regularization) transform microbiome and host omics count data into separate latent spaces, which are fused via a concatenation layer to form **Z**^*Join*^.

The two variants of DBayesCM are identical in encoder architecture, prior structure, and training procedure, and differ only in the geometry of the shared latent space and the corresponding mixture family, isolating the effect of latent geometry on clustering. DBayesCM-vMF constrains **Z**^*Join*^ to the unit hypersphere and clusters it with a von Mises-Fisher (vMF) mixture, so that samples are grouped by the relative direction of their multi-omic profile rather than its magnitude, a property suited to compositional data [16, 17]. For DBayesCM-vMF, the encoder outputs parameterize a vMF variational distribution with mean direction ***Z***_***µ***_ (normalized to unit length) and concentration ***Z***_***k***_ (exponentially transformed to ensure positivity) [17, 26]. DBayesCM-GMM uses a Euclidean latent and a Gaussian mixture [19]; its encoder outputs mean ***Z***_***ρ***_ and variance ***Z***_***σ***_, following the standard VAE formulation [27]. Full architectural specifications, including layer dimensions, activation functions, dropout rates, and the reparameterization used for each variational family, are provided in the Supplementary Material.

#### Nonparametric mixture models

DBayesCM-vMF expresses vMF mixture model with an infinite number of mixture components as follows:

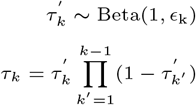

where *V* (.) denotes the vMF distribution, 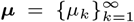 denotes a set of mean directions, and 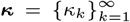 denotes a set of concentration parameters, ℐ_*s*_(.) is the modified Bessel function of the first kind of order s, 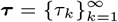 denotes a set of mixture coefficients, satisfying *τ*_*k*_ *>* 0 and 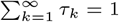.

The infinite mixture is constructed via a Dirichlet process prior using the stick-breaking representation [13], which enables the automatic determination of the effective number of clusters:

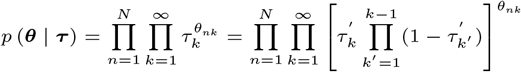

The cluster proportion *θ*_*nk*_ represents the posterior probability that sample *n*^*th*^ belongs to cluster *k*^*th*^, with the prior distribution

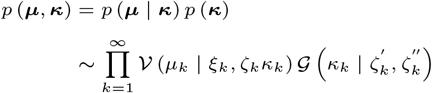

We introduce a vMF-Gamma distribution as the prior over ***µ*** and ***k*** as follows:

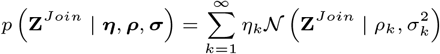

where *G*(.) denotes the gamma distribution, and *ϵ*_*k*_, *ξ*_*k*_, *ζ*_*k*_, 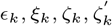 and 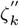 are hyperparameters of the *k*^*th*^ mixture component.

DBayesCM-GMM implements an infinite Gaussian mixture model on **Z**^*Join*^ which has been widely validated in clustering applications [19]:

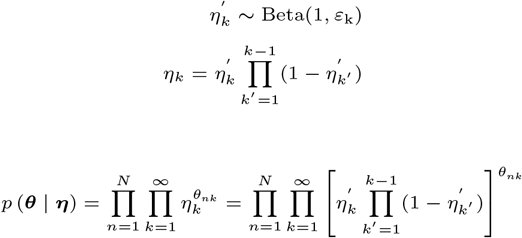

where *N* (.) denotes the normal distribution with mean *ρ*_*k*_ and variance 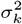. The DP prior is similarly constructed via stick breaking:

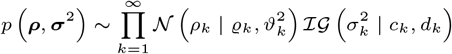

with conjugate Normal-Inverse-Gamma priors:

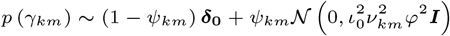

where ℐ*G*(.) denotes the Inverse-Gamma distribution, and *ε*_*k*_, *η*_*k*_, *ρ*_*k*_,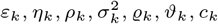, *ϱ*_*k*_, *ϑ*_*k*_, *c*_*k*_ and *d*_*k*_ are prior hyperparameters. Both variants enable probabilistic cluster assignments that quantify the uncertainty in cluster membership.

#### Feature selection

To identify the minimal set of discriminative features that characterize each cluster, we modeled cluster-specific embeddings *γ*_*km*_ (microbiome taxa) and *λ*_*ko*_ (host omics features) using spike-and-slab shrinkage priors [20]. This Bayesian variable selection approach probabilistically determines which features contribute meaningful information to cluster *k*^*th*^, automatically removing irrelevant features to enhance both the clustering performance and interpretability.

The spike-and-slab prior for microbiome embedding *γ*_*km*_ combines a point mass at zero (the “spike,” representing feature exclusion) with a continuous Normal distribution (the “slab,” representing feature inclusion):

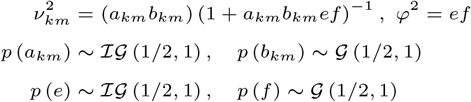

where the binary indicator *ψ*_*km*_ ∈ *{*0, 1*}* determines feature inclusion (*ψ*_*km*_ = 1 indicates taxonomic unit *m*^*th*^ is selected for cluster *k*^*th*^; *ψ*_*km*_ = 0 indicates exclusion), ***δ***_**0**_ denotes the Dirac delta function, and ***I*** is the identity matrix. The indicator follows a Bernoulli prior *p* (*ψ*_*km*_ | *ω*_*km*_) *∼* Bernoulli (*ω*_*km*_) with *ω*_*km*_ controlling the prior inclusion probability.

The slab component specifies a normal prior with a hierarchical variance structure that captures both feature-specific and global shrinkage. The slab variance decomposes into local shrinkage 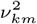 (controlling element-wise exclusion of taxonomic units irrelevant to cluster *k*^*th*^), global shrinkage *φ*^2^ (regulating overall sparsity across all features), and a known constant 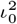 (set to stabilize posterior inference). Both shrinkage terms are constructed from auxiliary scalars through the products *a*_*km*_*b*_*km*_ and *ef*, where each auxiliary variable is assigned a conjugate Inverse-Gamma or Gamma prior. This scale-mixture parameterization induces heavy-tailed shrinkage priors on the variance components while retaining conditional conjugacy, enabling efficient posterior inference [20]:

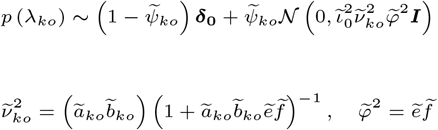

where *G*(.) and ℐ*G*(.) represent the gamma and inverse gamma distributions, respectively, and ***I*** is the identity matrix.

An identical spike-and-slab structure applies to the host omics embeddings *λ*_*ko*_:

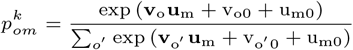

with analogous hyperpriors 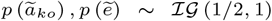,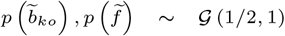, and an inclusion indicator 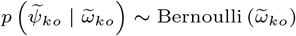. The posterior distributions of *ω*_*km*_ and 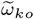 identify the minimal set of microbial taxa and host molecular features that jointly define each cluster, facilitating interpretation and biomarker discovery.

#### Prediction of co-occurrence relationships

To model the conditional probability relationships between selected microbial species and host omics features within clusters, we adapted VBayesMM (Variational Bayesian Microbiome Multiomics) [21], a Bayesian neural network approach that predicts host omics from microbial metagenomics. VBayesMM was applied to the subset of features identified as discriminative by the spike-and-slab feature selection of DBayesCM in the decoder step, thereby operating on a reduced feature space.

For a given cluster *k*^*th*^ with selected taxonomic units 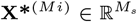and host omics features 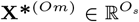(where *M*_*s*_ and *O*_*s*_ denote the number of selected features), VBayesMM models the conditional distribution using an encoder-decoder architecture. The encoder maps microbiome counts to a latent low-dimensional space of dimension *L*^*′*^ via embedding matrix 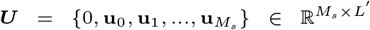 with prior *p* (***U*** ) *∼ N*(0, **Σ**_***U***_ ^2^ ). The decoder generates omics predictions through embedding matrix 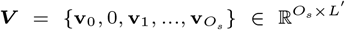with prior *p* (***V*** ) *∼ N* (0, **Σ**_***V***_ ^2^ ). For cluster *k*^*th*^, the conditional probability of observing host omics feature *o*^*th*^ given taxonomic unit *m*^*th*^ is

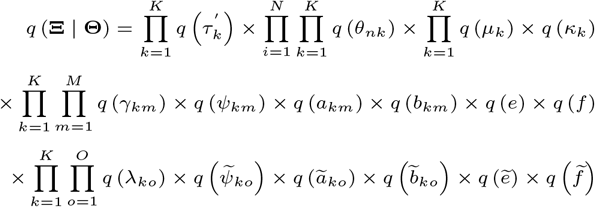

where *v*_*o*0_ and *u*_*m*0_ are the bias terms. For cluster *k*^*th*^, host omics features follow a multinomial distribution: 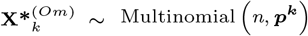. For each cluster and omics modality, VBayesMM outputs conditional log-probabilities log (***p***^***k***^ )revealing cluster-specific pathways by which the microbiome differentially regulates metabolomic, transcriptomic, and post-transcriptional programs. Mathematical explanations are provided in the Supplementary Material.

### Variational Bayesian approach

Exact posterior inference in DBayesCM is intractable because of the nonconjugate coupling between deep neural network encoders and infinite mixture models. Therefore, we employ the variational Bayesian (VB) method [13, 21], which approximates the intractable posterior with a tractable variational distribution optimized to minimize the Kullback-Leibler divergence from the true posterior.

#### Variational families

Given the observed omics datasets **X**^(*Mi*)^ and **X**^(*Om*)^, defines the complete parameter set **Ξ** comprising cluster assignments ***θ***, stick-breaking weights ***τ***^***′***^, vMF cluster parameters *{****µ, k****}*, microbiome embeddings *{****γ, ψ, a, b, e, f*** *}* and host omics embeddings 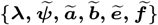. DBayesCM-GMM analogously defines **ψ** with stick-breaking weights ***η***^***′***^ and Gaussian cluster parameters *{****ρ, σ****}* replacing the vMF components while retaining identical embedding structures. We adopt mean-field variational inference, which assumes independence among latent variables in the variational distribution *q* (**Ξ** | **Θ**) (characterized by the hyperparameter **Θ**) and *q* (**ψ** | **Ω**) (characterized by **Ω**).

To render the infinite Dirichlet process computationally feasible, we employ truncated stick-breaking with a finite truncation level K [17, 19]. Critically, K serves as an upper bound rather than a model selection parameter: rather than fixing the number of clusters, the variational objective penalizes unnecessary components, and the stick-breaking weights of redundant components shrink toward zero during optimization. We therefore define the effective number of clusters as the number of components retaining non-negligible posterior mass at convergence, specifically, components to which at least one sample is assigned under the maximum-a-posteriori rule argmax_*k*_ *θ*_*nk*_. The factorized variational distribution for DBayesCM-vMF is with an analogous factorization for *q*(**ψ** | **Ω**) substiting Gaussian parameters *q* (*ρ*_*k*_), *q* (*σ*_*k*_) and 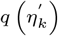 for the vMF components. Exponential-family variational distributions were selected to guarantee tractable expectations; modality-specific considerations are detailed in the Supplementary Material.

#### Evidence lower bound optimization

The VB approach maximizes the Evidence Lower Bound (ELBO) with respect to the variational parameters **Θ** (or **Ω**). The ELBO function of DBayesCM-vMF is expressed as follows:

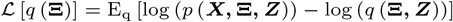

where ***X*** = *{****X***^*Mi*^, ***X***^*Om*^*}*, **Θ** includes the learnable parameters of both the fully connected neural networks and variational posteriors. The first term represents the expectation of the joint distribution with respect to the variational distribution, whereas the second term (negative entropy) penalizes overly concentrated posteriors. The gradient with respect to the variational parameters is

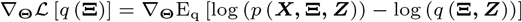

DBayesCM-GMM employs the analogous ELBO ℒ [*q* (**ψ**)] and gradient *∇*_**Ω**_ ℒ [*q* (**ψ**)] with Normal latent space assumptions. Since the variational distributions *q* (**Ξ**) and *q* (**ψ**) are reparameterized into differentiable forms (via the trick for continuous variables and Gumbel-softmax for discrete indicators), we can utilize the stochastic gradient approach [28] to optimize the ℒ [*q* (**Ξ**)] and ℒ [*q* (**ψ**)]. The mathematical details of the gradients *∇*_**Θ**_ ℒ [*q* (**Ξ**)] and *∇*_**Ω**_ ℒ [*q* (**ψ**)] with respect to the variational parameters are provided in the Supplementary Material.

#### Hyperparameter initialization

In all our experiments, we initialized the truncation level for the infinite mixture models at 10 components, which the variational inference procedure automatically refines by collapsing the unnecessary clusters. The stick-breaking concentration parameters were set to *ϵ* = *ε* = 0.1 [13, 29]. To address the selection of microbial species and host omics features, we set the the initial value of the hyperparameter ***ω*** to 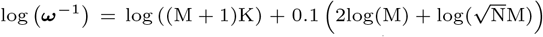 and 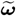 to 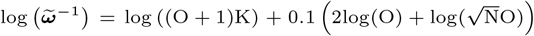 [21, 20]. Owing to significant differences in the number of microbial features (M), host omics features (O), and samples (N) across our datasets, the actual values of ***ω*** and 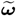 differ for each dataset. This formulation imposes stronger sparsity priors in datasets with larger M or O relative to N, automatically adapting to each dataset’s feature-to-sample ratio and mitigating the overfitting. For DBayesCM-vMF, vMF mean directions ***ξ*** were initialized to the data centroid (the empirical mean of the latent representations), concentration scaling ***ζ*** = 0.01, and Gamma hyperparameters (***a, b***) = (1.0, 0.1) [16, 17]. DBayesCM-GMM sets the initial values of the hyperparameters ***ϱ, ϑ, c, d*** to 1.0. A comprehensive explanation of the initial values of the hyperparameters of all priors is provided in the Supplementary Materials.

### Benchmarking and evaluation

**MOFA+** decomposes multi-omics data into latent factors capturing shared and modality-specific variation; it does not directly cluster. We ran MOFA+ (R package v1.20.2) with default parameters, extracted 15 latent factors, and applied *k*-means clustering with *K* chosen by the Bayesian information criterion (BIC). **iCluster** performs joint latent-variable modelling via generalized linear models with lasso penalization, assuming a Poisson distribution for counts and requiring a pre-specified *K*. We ran iCluster (R package v1.46.0) with default parameters and selected *K* via BIC. **MultiVI** is a VAE-based generative model integrating paired modalities through modality-specific encoders and a shared latent space. We ran MultiVI (scvi-tools v1.3.3) with default parameters and applied *k*-means clustering on the latent representation with *K* chosen by BIC. **SnapCCESS** generates multiple embeddings via snapshot-ensemble VAEs with cyclic learning-rate scheduling; consensus clustering was derived from *k*-means outputs across snapshots, with *K* chosen by BIC.

The clustering performance was assessed using the Adjusted Rand Index (ARI), which measures the agreement between the predicted cluster assignments *C* = *{C*_1_, …, *C*_*k*_*}* and true labels *T* = *{T*_1_, …, *T*_*m*_*}* while correcting for chance:

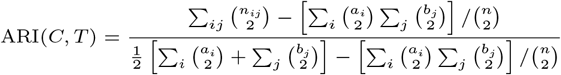

where *n*_*ij*_ is the number of samples in cluster *C*_*i*_ and true class *T*_*j*_, *a*_*i*_ = Σ _*j*_*n*_*ij*_, *b*_*j*_ = Σ _*i*_*n*_*ij*_, and *n* is the total sample size. The ARI ranges from 0 (random agreement) to 1 (perfect agreement).

The prediction accuracy for host omics features from taxonomic units was evaluated using the Symmetric Mean Absolute Percentage Error (SMAPE):

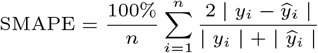

where *y*_*i*_ is the observed host omics feature value (metabolite abundance, gene expression, or miRNA expression), and *y*_*i*_ is the predicted value from the taxonomic unit input for sample i across n test samples. SMAPE ranges from 0% (perfect prediction) to 100% (maximum error), is symmetric for over- and under-predictions, and avoids instability near zero values.

### Datasets

We evaluated DBayesCM on four published multimodal omics datasets spanning human cancers and mouse disease models, which were selected to represent diverse tissue types and cancer stages. Dataset A comprises fecal microbiome (623 taxonomic units) and metabolite profiles (450 features) from colorectal cancer (CRC) patients stratified by disease stage (I/II, III/IV) and healthy controls [22]. Datasets B and C derive from the integrated TCGA-TCMA cancer microbiome databases [24, 25]. Dataset B profiles the blood microbiome (2,023 taxa), tumor RNA-seq (868 genes), and miRNA expression (956 features) from patients with kidney renal clear cell carcinoma (KIRC) across stages I, III, and IV. Dataset C profiles the breast tumor microbiome (3,584 taxa), RNA-seq (2,135 genes), and miRNA expression (823 features) from patients with breast invasive carcinoma (BRCA) across stages I, IIB, and IIIA. Dataset D comprises gut microbiome (4,690 taxa) and metabolite profiles (1,710 features) from a murine obstructive sleep apnea (OSA) model comparing intermittent hypoxia-hypercapnia (IHH) versus control conditions [23]. We note that tumor- and blood-derived microbiome profiles from TCGA-based resources have been reported to be substantially affected by contamination and batch effects, particularly in low-biomass tissues [30]. Datasets B and C are therefore used here to demonstrate that DBayesCM operates on heterogeneous multi-modal cancer data structures (microbiome, miRNA, and RNA-seq). For each dataset, we randomly partitioned the samples into training (80%) and test (20%) sets, maintaining class balance across disease stages or conditions.

## Results

### DBayesCM recovers known cluster structure under controlled conditions

To evaluate whether a hyperspherical latent space offers an advantage under controlled, ground-truth conditions, we simulated N=360 compositional count profiles (120 features) from four balanced groups, with per-sample sequencing depth drawn uniformly from 4,000–12,000 reads to introduce realistic magnitude variation unrelated to group identity [31]. Both DBayesCM variants were fit under matched encoder architecture and training conditions, differing only in output of mixture family, using the same finite truncation level K = 10 applied throughout this study (Fig. 2). From this shared truncation, DBayesCM-vMF automatically pruned to exactly four effective clusters, matching the ground truth (ARI = 0.991; Fig. 2d), whereas DBayesCM-GMM retained five effective clusters under the identical Dirichlet-process prior, splitting one true group into two (ARI = 0.897; Fig. 2b). The two geometries also differ in how compactly they summarize a recovered cluster: a vMF component reduces to a single mean direction and concentration, so the separation between any two clusters is captured by one interpretable, rotation-invariant quantity, the angle between their mean directions, whereas a Gaussian component requires location, orientation, and shape (Fig. 2b, ellipsoids) with no equivalent single measure.

**Fig. 2.**
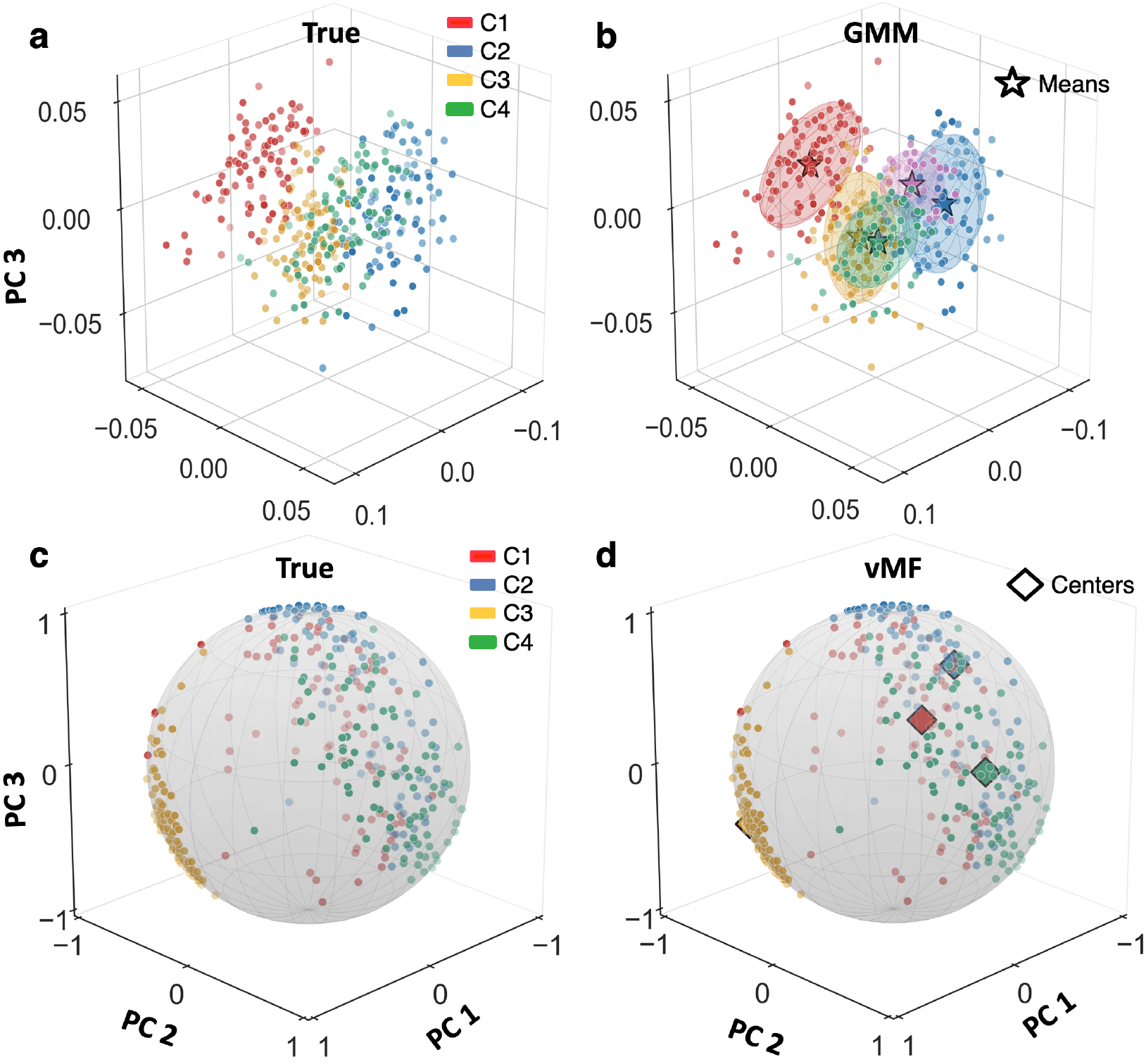
A hyperspherical (vMF) mixture recovers the latent structure and number of clusters under controlled conditions. Compositional count profiles (360 samples, 120 features) were simulated from four balanced ground-truth groups (C1-C4). Top row: the shared latent space of DBayesCM-GMM (standard Gaussian encoder, Euclidean latent).(a) Ground-truth cluster assignments. (b) recovered Gaussian mixture components shown as coverage ellipsoids with their means (stars). Bottom row: the shared latent space of DBayesCM-vMF (hyperspherical encoder, unit-sphere latent). (c) the same ground-truth assignments on the unit hypersphere. (d) recovered von Mises-Fisher mixture components with their mean directions marked as diamonds. Both variants were fit under matched encoder architecture and training conditions from the same finite truncation level K = 10.

All panels show a three-dimensional PCA projection of the respective latent space and are provided for visualization; all angular quantities were computed in the full latent space. There, the four recovered vMF mean directions lie within 14.2 − 22.6°of one another (mean 19.8°), indicating that the recovered clusters occupy a comparatively narrow angular neighbourhood. This narrow true gap shows that DBayesCM-vMF distinguishes groups primarily through the full joint likelihood, not through wide angular separation of cluster archetypes.

Having established that the directional formulation recovers known structure and automatically identifies the correct number of clusters under controlled conditions, we next apply DBayesCM to large-scale public datasets with known ground truths and benchmark its clustering accuracy and uncertainty quantification against SnapCCESS [7], MultiVI [8], MOFA+ [10], and iCluster [11].

### Benchmarking clustering performance across diseases and data modalities

We first evaluated the clustering performance of DBayesCM on four multiomics microbiome datasets by benchmarking it against four established methods. While iCluster performs intrinsic sample clustering, MOFA+ detects latent factors, and SnapCCESS and MultiVI learn embeddings. For methods that did not produce direct cluster assignments, we applied k-means consensus clustering to their factor or embedding matrices to enable a fair comparison. DBayesCM consistently achieved superior clustering accuracy across all cancer types, non-cancer disease stages, and data modality combinations (Fig. 3).

**Fig. 3.**
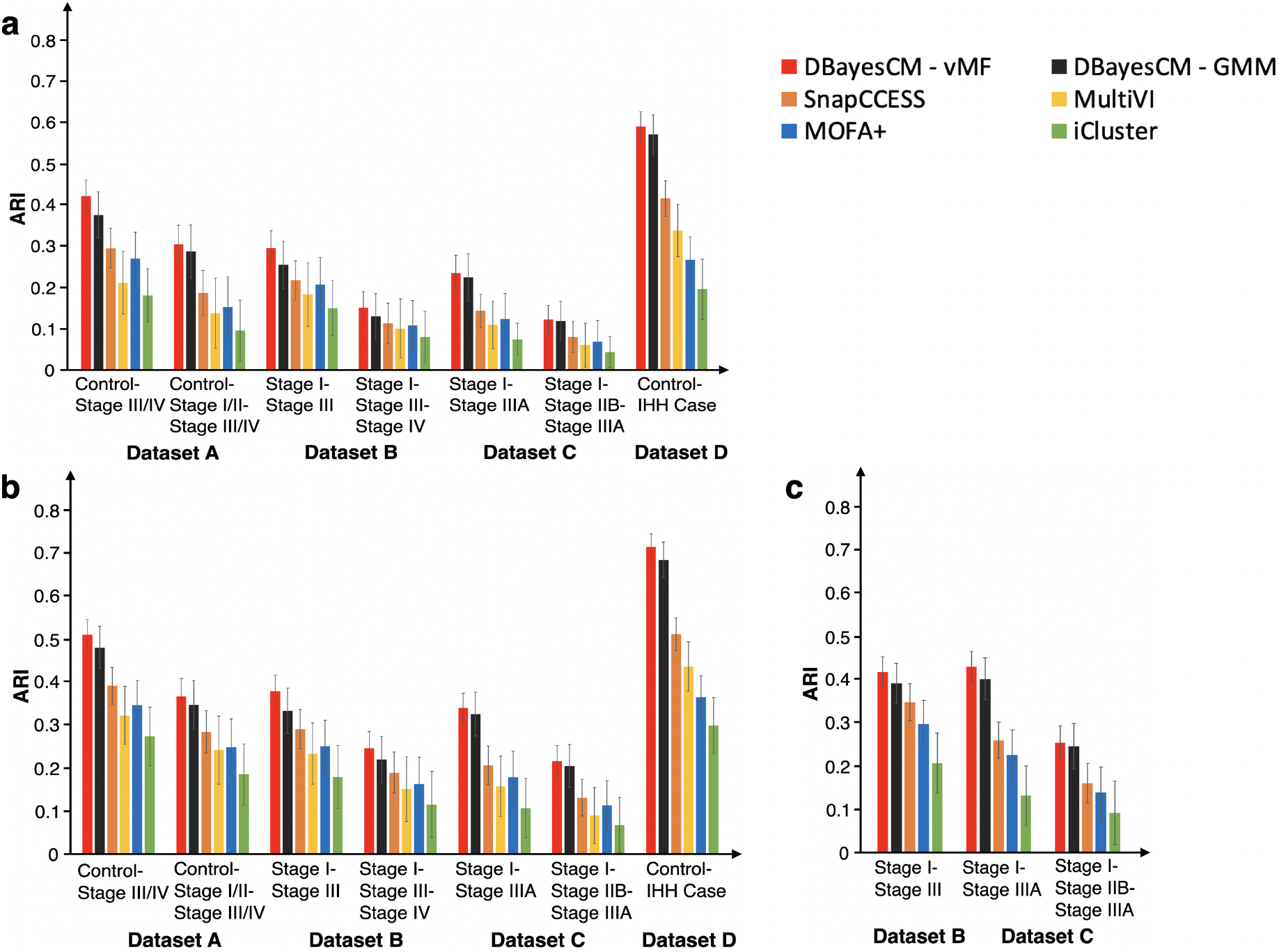
Clustering performance comparison across diseases and data modalities. Adjusted rand index (ARI) values for DBayesCM and baseline methods across the four real-world datasets. Dataset A: colon cancer (CRC) with stage I/II, stage III/IV, and control samples. Dataset B: Kidney cancer (KIRC) with stages I, III, and IV. Dataset C: breast cancer (BRCA) samples of stages I, IIB, and IIIA. Dataset D: obstructive sleep apnea (OSA) with IHH in cases and controls. DBayesCM was evaluated using two mixture models: von Mises-Fisher mixture model (DBayesCM-vMF) and Gaussian mixture model (DBayesCM-GMM) for the clustering process. **(a)** microbiome data only. **(b)** microbiome-metabolite for datasets A and D, microbiome-miRNA for datasets B and C. **(c)** microbiome-miRNA-RNAseq for datasets B and C data. Note: All algorithms were run on a personal computer (Intel Xeon Gold 6230 Processor 2.10 GHz × 2, 40 cores, and NVIDIA Quadro GV100) under Ubuntu 24.04.3 LTS.

For microbiome data alone (Fig. 3a), DBayesCM-vMF demonstrated substantial performance advantages across diverse clinical conditions. For colon cancer (Dataset A), DBayesCM-vMF achieved ARI values ranging from 0.30 (SD = *±*0.046) (three-group clustering) to 0.42 (SD = *±*0.040) (two-group clustering), outperforming the SnapCCESS method (0.18 *±*0.055 and 0.29 *±*0.048, respectively) by margins of 0.12-0.13. Inter-stage comparisons of kidney cancer (Dataset B) and breast cancer (Dataset C) showed modest performance, indicating that microbiome signatures alone provide limited discriminative power between adjacent cancer stages. In contrast, distinguishing severe disease from controls showed markedly higher accuracy: Control-IHH cases in obstructive sleep apnea (Dataset D) achieved ARI values of 0.58 (SD = *±*0.036) for DBayesCM-vMF versus 0.42 (SD = *±*0.043) for SnapCCESS.

The integration of the microbiome with host omics data enhanced the clustering performance across all methods, with DBayesCM exhibiting the most pronounced improvements (Fig. 3b). For colon cancer with microbiome-metabolite integration (Dataset A), DBayesCM-vMF achieved ARI values from 0.37 (SD = *±*0.042) for two-group clustering, improving over microbiome-only analysis in both cases. To confirm that these values reflect genuine structure rather than chance agreement, we tested each Dataset A against a permutation null (1000 permutations; Supplementary Table S1). All four comparisons were significant, with empirical *p ≤* 0.001; every null distribution was centred near zero.

The performance gap between DBayesCM and competing methods widened with data integration: whereas SnapCCESS improved ARI values from 0.29 (SD = *±*0.048) to 0.40 (SD = *±*0.043), DBayesCM-vMF increased from 0.42 to 0.51 for two-group clustering, demonstrating a superior ability to extract complementary information from heterogeneous data types. This advantage was most striking for obstructive sleep apnea (Dataset D), where DBayesCM-vMF reached ARI value of 0.72 (SD = *±*0.030) compared to 0.52 (SD = *±*0.038) for SnapCCESS and 0.37 (SD = *±*0.051) for MOFA+. For kidney cancer with microbiome miRNA integration (Dataset B), DBayesCM-vMF maintained superior performance with ARI values of 0.25-0.38 across comparisons. Breast cancer three-stage discrimination (Dataset C) proved to be the most challenging across all methods, with DBayesCM-vMF reaching an ARI value of 0.22 (SD = *±*0.036) compared to 0.13 (SD = *±*0.043) for the SnapCCESS method, reflecting the high molecular heterogeneity within breast cancer groups. The permutation null test applied to kidney (Dataset B) and breast cancer (Dataset C) yielded significant results across all inter-stage comparisons (Supplementary Table S1).

The addition of RNA-seq data to microbiome miRNA integration further enhanced cluster discrimination (Fig. 3c). For kidney cancer, DBayesCM-vMF reached an ARI value of 0.42 (SD = *±*0.035) versus 0.35 (SD = *±*0.043) for SnapCCESS and 0.30 (SD = *±*0.056) for MOFA+. For breast cancer three-stage discrimination, DBayesCM-vMF achieved an ARI value of 0.43 (SD = *±*0.036), outperforming SnapCCESS (0.26 *±*0.042) and MOFA+ (0.23 *±*0.058). This suggests that miRNA dysregulation provides critical discriminative information for these cancer types. Both DBayesCM variants consistently ranked first and second for all the conditions tested. The consistent top-two ranking across diverse cancer types (CRC, KIRC, and BRCA) and a non-cancer disease (OSA) demonstrated a robust generalization capability. This suggests that the Bayesian framework of DBayesCM is particularly effective for capturing complex interactions in heterogeneous multiomics data, where traditional methods struggle.

### DBayesCM provides probabilistic cluster assignments and hyperspherical cluster geometry

A critical limitation of existing multiomics clustering methods is their inability to quantify the confidence in cluster assignments. DBayesCM addresses this through its Bayesian framework, which provides posterior probabilities for each sample’s cluster membership, enabling the identification of both confident and ambiguous assignments. The following analyses and illustrative results are presented for dataset A (colorectal cancer), the comprehensively characterized cohort in this study (Figs. 4 and 5).

**Fig. 4.**
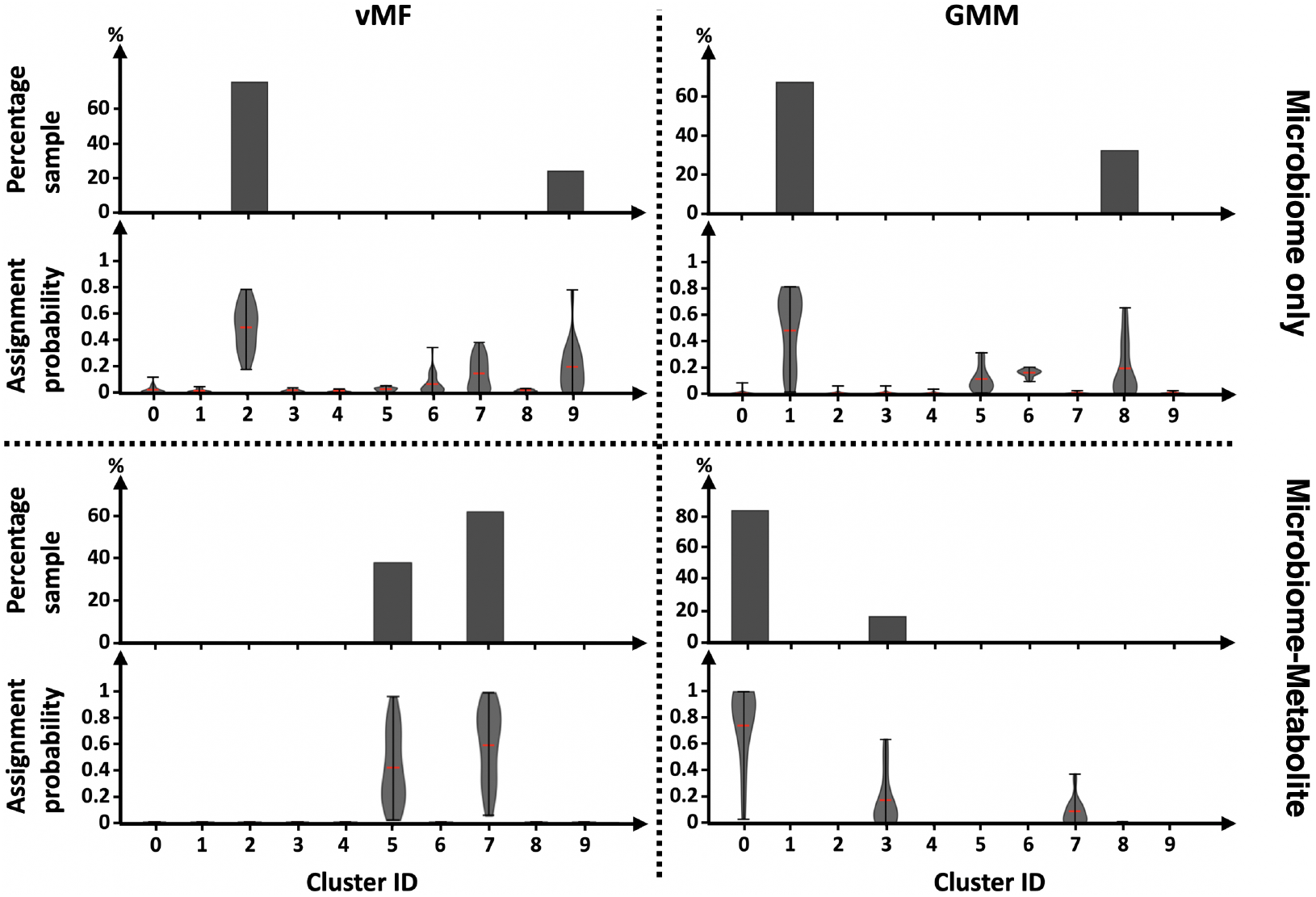
Probabilistic cluster discovery with uncertainty quantification for two-group structure of dataset. **A.** Top: Percentage of samples assigned to each cluster (0-9). Bottom: Posterior assignment probabilities (violin plots; red lines indicate means) quantifying confidence for each cluster. vMF (von Mises-Fisher mixture model) and GMM (Gaussian mixture model) clustering applied to microbiome-only (upper row) versus integrated microbiome-metabolite data (lower row).

**Fig. 5.**
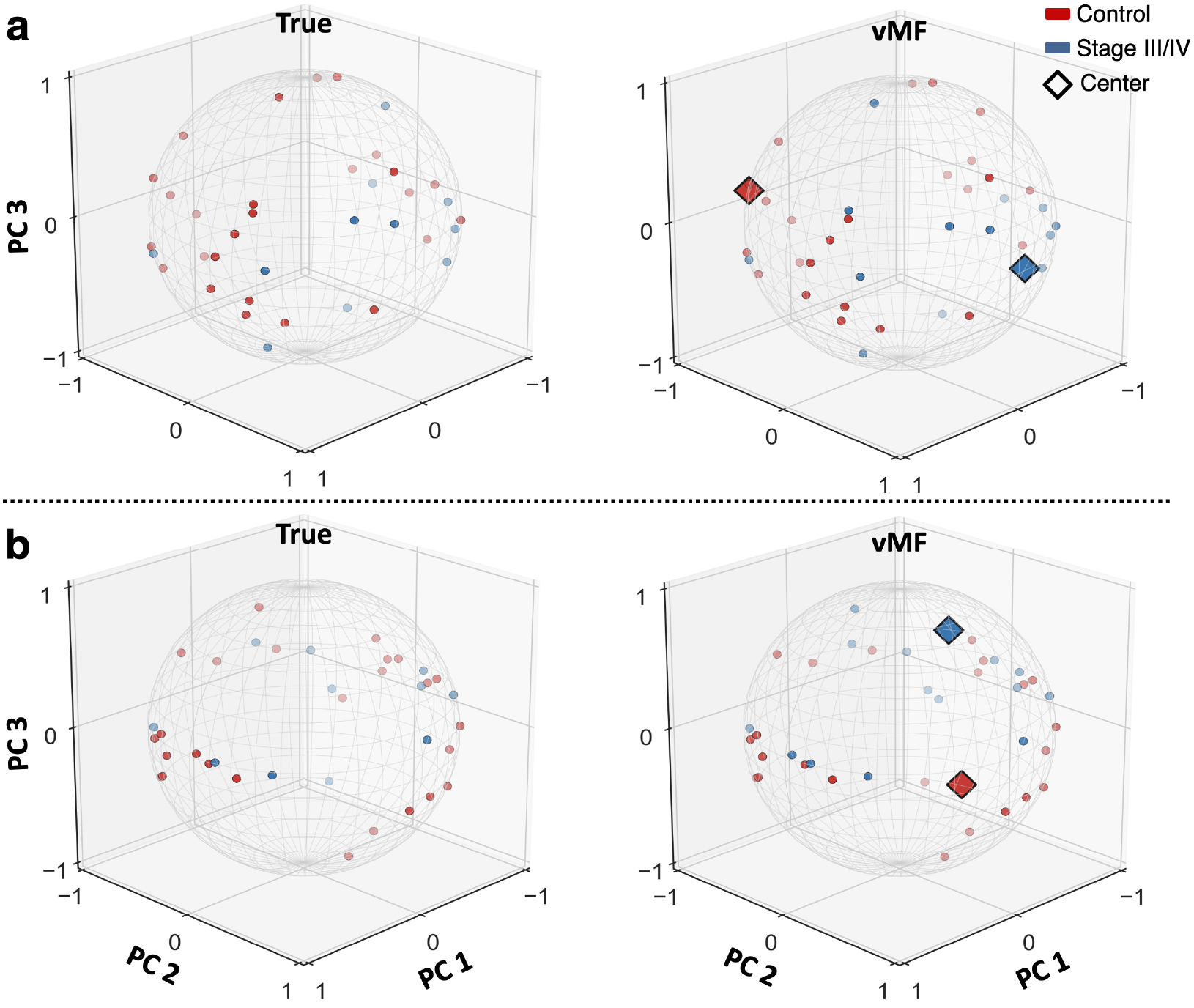
Visualization of multi-omics integration on the recovered cluster structure for the two-group comparison of dataset. **A.** Sample positions are shown as a three-dimensional PCA projection (PC1–PC3) of the L-dimensional unit-hypersphere latent space (L = 64). “True” panels (left) show the ground-truth partition; “vMF” panels (right) show the same samples with the two highest-weight von Mises–Fisher mixture components overlaid, their mean directions marked as diamonds (“Center”). Red: healthy controls; blue: Stage III/IV cancer. **(a)** microbiome data only. **(b)** microbiome-metabolite data.

Microbiome data alone produces broad posterior uncertainty (Fig. 4, top row). For the two-group comparison (control and stage III/IV), DBayesCM-vMF distributed posterior mass across clusters 2 (*∼* 75% of samples, mean *∼* 0.50) and 9 (*∼* 25% of samples, mean *∼* 0.30), with additional probability in clusters 0, 5, 6, and 7. DBayesCM-GMM showed similar uncertainty patterns across clusters 1 (*∼* 68% of samples, mean *∼* 0.75) and 8 (*∼* 32% of samples, mean *∼* 0.28). This proliferation indicates that the model cannot confidently discriminate between groups given the limited single-modality data.

The addition of metabolite data produced a striking transformation in the posterior distributions (Fig. 4, bottom row). For the two-group comparison, DBayesCM-vMF achieved clean assignments with violin plots appearing at only two cluster positions (clusters 5 and 7), precisely matching the true structure. Cluster 7 showed a narrow, high-confidence distribution (*∼* 62% of samples, mean *∼* 0.60), while the absence of probability mass at other positions demonstrates the model confidently identifies biological structure without spurious cluster proliferation. DBayesCM-GMM showed improvement but retained residual uncertainty across multiple clusters (0, 3, and 7), with cluster 0 achieving high confidence (mean *∼* 0.73), while cluster 3 showed ambiguity (mean *∼* 0.18). For the more complex three-group discrimination (control-stage I/II-stage III/IV), DBayesCM-vMF identified exactly three clusters (2, 3, and 5) with probability mass concentrated at these positions only (Supplementary Fig. S1). DBayesCM-GMM distributed probability across four cluster positions despite making three-cluster assignments (0, 4, and 6), with residual probability at a spurious cluster 9 (mean *∼* 0.10), indicating lingering model uncertainty about whether the true structure contains three or four clusters. This pattern demonstrates how directional clustering on the hypersphere provides a more decisive cluster resolution for high-dimensional omics data.

The hyperspherical projection in Fig. 5 illustrates a property specific to directional clustering: because the encoder maps each sample to a unit-length latent vector, position on the sphere encodes the relative shape of a sample’s multi-omic profile, and cluster membership is governed by angular proximity to a discovered mean direction rather than Euclidean distance to a centroid. The two center markers correspond to the model’s estimated mean directions for the Control and Stage III/IV groups. In the full 64-dimensional latent, these two directions are separated by 7.6°with microbiome data alone (Fig. 5a) and 4.5°(Fig. 5b) with microbiome and metabolite data combined. A separation of a few degrees indicates that the two clusters share most of their profile and are distinguished along a narrow, disease-specific axis rather than occupying broadly separated regions of the latent space. This regime is resolved primarily by direction rather than magnitude, which suits the hyperspherical formulation. Because the encoder output is unit-normalized, variation in overall abundance or sequencing depth, unrelated to disease state, is removed from the clustering signal by construction, and the uniform prior on the sphere imposes no shrinkage toward a shared origin.

The contrast is visible on the same dataset: in the Euclidean latent of DBayesCM-GMM, encoded samples are compressed into a small region near the origin and the two recovered Gaussian components overlap substantially (Supplementary Fig. S2), whereas the hyperspherical representation distributes the same samples across the sphere with better-separated mean directions (Fig. 5). This parallels the controlled simulation (Fig. 2), in which the directional formulation recovered the correct number of components while the Gaussian variant did not. Consistent with this picture, metabolite integration is accompanied by a collapse in the number of mixture components carrying non-trivial posterior mass, from four with microbiome data alone to exactly two with metabolite data added (Fig. 4), coinciding with the improvement in clustering accuracy (ARI = 0.42 to 0.51). All angular quantities reported here were computed in the full 64-dimensional latent space; the three-dimensional projection in Fig. 5 is provided for visualization only, and individual sample positions in it are approximate.

### DBayesCM identifies interpretable microbiome-host omics features with quantified selection confidence

DBayesCM’s spike-and-slab approach identifies discriminative features while quantifying selection confidence. This approach assigns each feature a posterior selection probability, distinguishing genuinely discriminative variables from the noise. DBayesCM-vMF identified distinct high-confidence features (selection probability *>* 0.6) separated from ambiguous features (black points) (Fig. 6a), with bimodal probability distributions demonstrating appropriate uncertainty quantification rather than forced binary selections. DBayesCM-GMM showed comparable feature-selection patterns (Supplementary Fig. S3).

**Fig. 6.**
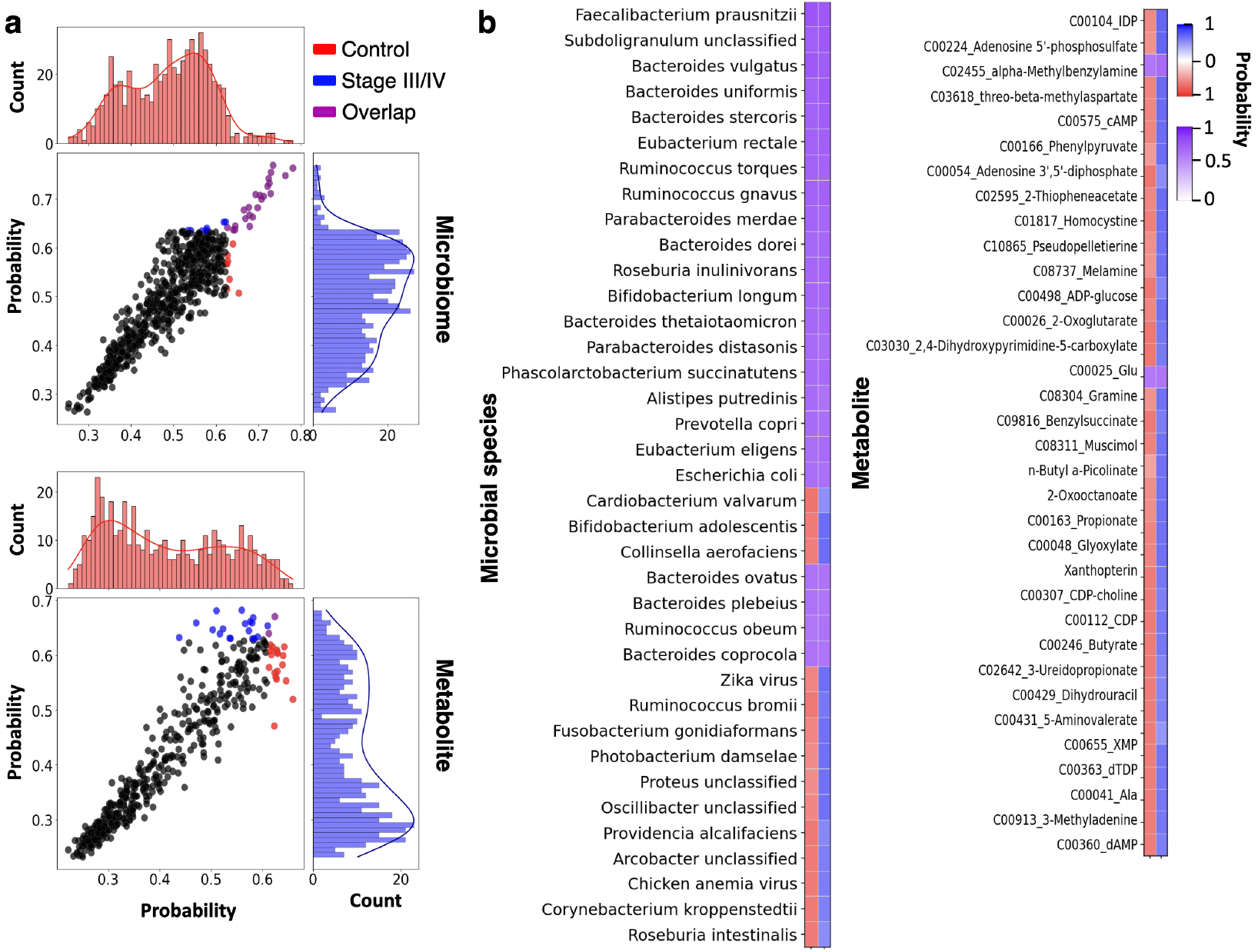
DBayesCM-vMF identifies discriminative microbiome-metabolite features with quantified selection confidence for dataset A. **(a)**Scatter plots and histograms of spike-and-slab posterior probabilities for microbiome and metabolite. **(b)** Heatmaps of top 30 microbial species (left) and top 20 metabolites (right) ranked by selection probability.

The bacterial taxa receiving the highest selection probabilities recapitulate well-established features of the colorectal cancer-associated gut microbiome (Fig. 6b left). *Faecalibacterium prausnitzii* achieved highest selection probability, consistent with its role as a major butyrate-producing commensal that significantly reduces aberrant crypt foci formation and suppresses HCT116 colorectal cancer cell proliferation in a time and dose-dependent manner [32], and is consistently depleted in CRC patients compared to healthy controls [22, 33]. Several other strongly selected taxa, including *Subdoligranulum, Roseburia*, and *Eubacterium* species, belong to the same guild of butyrate-producing Firmicutes that is characteristically reduced in colorectal cancer [34].

Metabolite selection identified multiple dysregulated pathways (Fig. 6b right). cAMP selection reflects aberrant cellular signaling, exhibiting context-dependent roles in CRC growth regulation and malignant progression through PKA-CREB pathways, with altered cAMP transport and turnover in CRC compared to healthy colonic mucosa [35]. The selection of adenosine 5^*′*^-phosphosulfate (APS) reflects the altered sulfate activation. APS is an intermediate in 3^*′*^-phosphoadenosine 5^*′*^-phosphosulfate (PAPS) biosynthesis, which is differentially expressed between metastatic and non-metastatic colon carcinoma cells and has recently been identified as a diagnostic biomarker in colon adenocarcinoma [36].

### DBayesCM reveals the co-occurrence relationships between the most informative microbial species and host omics features

To quantify the probabilistic relationships between the top 30 selected microbial species and top 20 selected metabolites, we applied Bayesian neural network inference from our VBayesMM package [21] to estimate conditional log-probabilities between these features for the stage III/IV and control groups. Both DBayesCM variants achieved mid-range symmetric mean absolute percentage error (SMAPE) values (*∼* 40% for stage III/IV, *∼* 39% for controls) when predicting metabolite profiles from microbial composition (Supplementary Fig. S4). Figure 7 displays the hierarchical clustering of conditional probabilities from DBayesCM-vMF, where teal and brown denote conditional associations of microbial species and metabolite that are stronger and weaker than average, respectively. DBayesCM-GMM exhibited comparable patterns (Supplementary Figure S5).

**Fig. 7.**
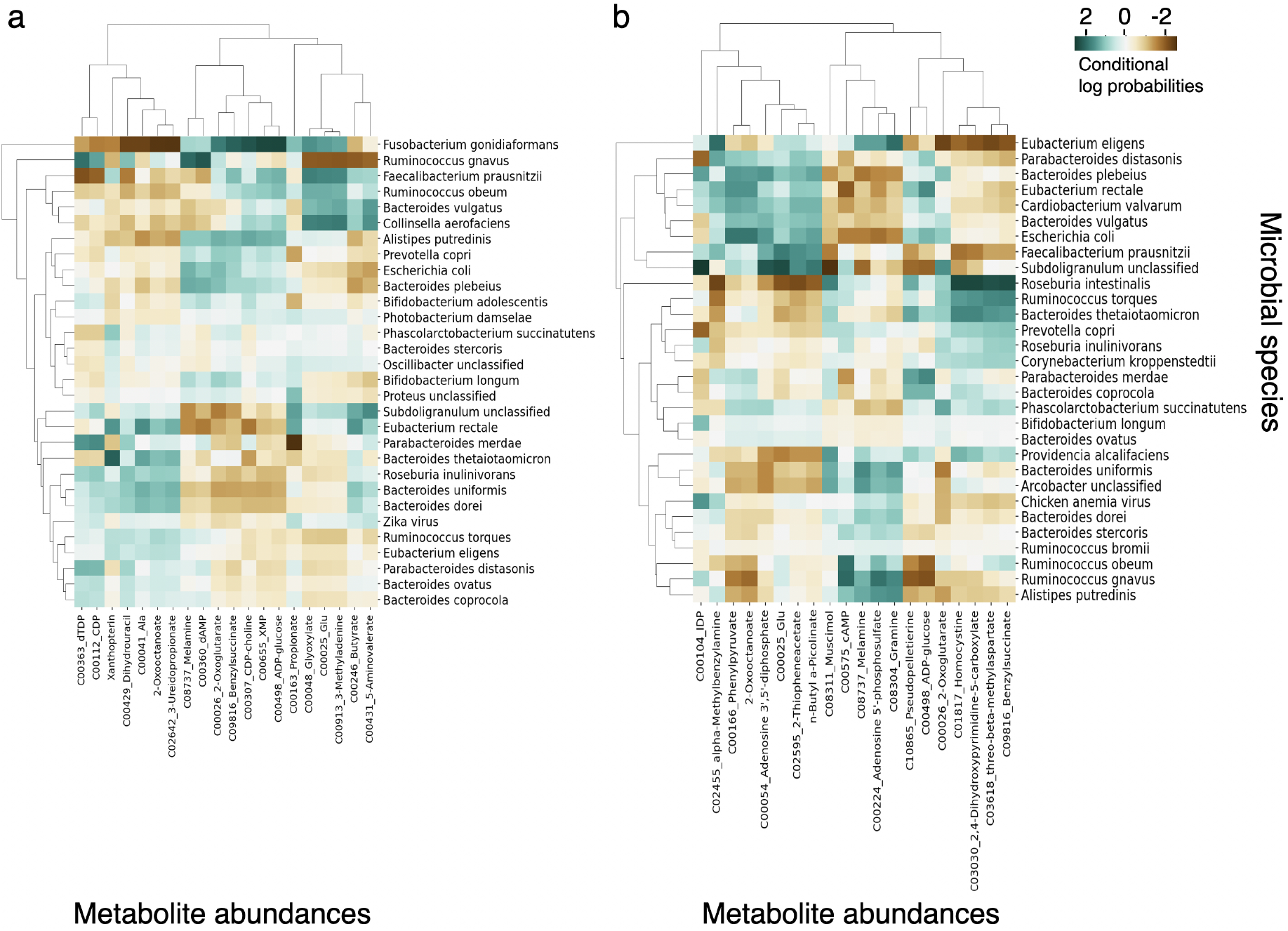
Heat map of the estimated conditional log probabilities of DBayesCM-vMF for the selected microbial species and metabolite abundances in dataset A. Individual metabolites and microbiomes were hierarchically clustered (Ward’s method) using Euclidean distance. (a) stage III/IV. (b) control group.

In stage III/IV (Fig. 7a), *Bacteroides* and *Clostridium* species exhibit positive conditional probabilities with secondary bile acids (deoxycholic acid, lithocholic acid). Conversely, *Fusobacterium* species show negative conditional probabilities ((indicated in brown)) with short-chain fatty acids (SCFAs). The control samples revealed different conditional structures (Fig. 7b). Butyrate-producing bacteria (*Faecalibacterium prausnitzii, Subdoligranulum*, and *Roseburia*) demonstrated positive conditional probabilities with SCFA metabolites and other beneficial compounds, capturing cooperative cross-feeding, where acetate utilization drives butyrate production [37, 38]. This Bayesian quantification distinguishes pathogenic metabolite suppression from protective metabolite cooperation, thereby providing hypotheses for precision oncology follow-up.

## Discussion

We introduce DBayesCM and apply it to microbiome multiomics datasets spanning colorectal, kidney, and breast cancers, as well as obstructive sleep apnea. DBayesCM has three key components. First, it integrates heterogeneous multiomics data (microbiome, metabolomics, RNA-seq, and miRNA) for cluster discovery using separate encoders for each modality combined through a joint latent space. Second, it performs automatic cluster discovery using infinite von Mises-Fisher mixture models with Dirichlet process priors, providing uncertainty quantification over cluster assignments and numbers. Third, it identifies a limited number of discriminative features through spike-and-slab shrinkage priors and predicts co-occurrence relationships between microbiome and omics features using Bayesian neural networks.

DBayesCM addresses several critical challenges in the integration of multiomics data. First, it provides full posterior distributions over cluster assignments rather than point estimates, enabling the identification of low-confidence assignments that require additional clinical investigation. Second, unlike competing methods that require pre-specified cluster numbers, DBayesCM automatically determines the optimal cluster count using Dirichlet process priors. For the two-group structure of the colon cancer study, DBayesCM-vMF concentrated probability at exactly two clusters (with metabolite integration), while for three-group comparisons, it identified precisely three clusters with sparse assignments, whereas DBayesCM-GMM showed residual uncertainty across additional positions. The hypersphere geometry of the vMF formulation appears to be particularly effective for high-dimensional omics data. Third, the spike-and-slab approach quantifies feature selection confidence, distinguishing a core set of influential omics features from noise through posterior selection probabilities. Fourth, DBayesCM estimates conditional probabilities to facilitate the discovery of probabilistic relationships between host and the microbiome’s community structure and function, offering valuable insights for understanding their impact on human health and cancer.

DBayesCM incorporates Bayesian deep learning, which is specifically designed for uncertainty-aware multiomics integration in precision oncology. This capability is particularly important, given the clinical requirement to distinguish confident predictions from ambiguous assignments. DBayesCM (both vMF and GMM variants) consistently outperformed the existing methods in terms of clustering accuracy and explicitly identified samples with low posterior probabilities. In addition, DBayesCM revealed that integrating the microbiome with host omics substantially reduces prediction uncertainty by capturing complementary biological signals. Microbiome composition reflects the microbial community structure, whereas host omics (metabolomics, RNA-seq, and miRNA) quantify functional consequences and systemic responses, enabling more definitive patient stratification.

We note several limitations that motivate future developments. First, DBayesCM currently employs Dirichlet-multinomial distributions optimized for count data but cannot handle continuous intensity measurements, such as proteomics, which exhibit log-normal or heavy-tailed distributions requiring alternative likelihood formulations [39, 40, 41]. Second, while DBayesCM integrates three omics modalities for clustering, it predicts only pairwise relationships between the microbiome and individual host omics layers, which limits biological interpretation; extending to higher-order interactions across multiple modalities (for example, microbiome-metabolite-gene regulatory networks) would yield richer explanations but would require more advanced graphical modeling approaches [42, 43, 44]. Third, current validations employ bulk tissue multiomics data with complex, overlapping clusters. Although DBayesCM can be applied to single-cell microbiome data, it faces challenges due to increased clustering complexity when resolving 10-30 distinct cell types [45, 46, 47, 48]. Fourth, although variational inference enables tractable computation for current count-based modalities, scaling to comprehensive multi-omics (such as single-cell transcriptomics [49, 50]) encounters computational bottlenecks owing to combinatorial growth in latent space dimensionality and posterior complexity.

## Key Points

- DBayesCM is a deep Bayesian nonparametric approach that, within a single model, infers the number of clusters from the data, quantifies uncertainty in each sample’s cluster assignment, and selects discriminative features with posterior inclusion probabilities.
- Two variants that are identical except for the geometry of the shared latent space isolate the effect of latent geometry on clustering: DBayesCM-vMF constrains the latent to a unit hypersphere with a von Mises-Fisher mixture, while DBayesCM-GMM uses a Euclidean latent with a Gaussian mixture.
- On simulated data, the hyperspherical variant recovered the correct number of clusters, whereas the Euclidean variant over-segmented. Across four microbiome multi-omics cohorts, DBayesCM ranked consistently among the current methods, with recovered clusters significant against a label-permutation null.
- Integrating host omics with microbiome profiles improved clustering accuracy and reduced assignment ambiguity, and the estimated probabilistic co-occurrence between selected microbial species and host omics features provides interpretable, hypothesis-generating summaries of microbiome-host associations.

## Code availability

The software DBayesCM is available at https://github.com/tungtokyo1108/DBayesCM

## Author contributions

**Tung Dang**: Conceptualization, Methodology, Software, Validation, Formal analysis, Visualization, Writing – original draft, Writing – review & editing. **Artem Lysenko**: Conceptualization, Methodology, Investigation, Validation, Supervision, Writing – review & editing. **Tatsuhiko Tsunoda**: Conceptualization, Methodology, Funding acquisition, Project administration, Writing – review & editing.

## Competing interests

No competing interest is declared.

## Funding

This work was partly supported by JSPS KAKENHI Grant Number JP20H03240, JSPS KAKENHI Grant Number JP24K15175 and JST CREST Grant Number JPMJCR2231, Japan.

## Notes

### Competing Interest Statement

The authors have declared no competing interest.

## References

1. Vancheswaran Gopalakrishnan, Beth A Helmink, Christine N Spencer, Alexandre Reuben, and Jennifer A Wargo. The influence of the gut microbiome on cancer, immunity, and cancer immunotherapy. Cancer cell, 33(4):570–580, 2018.

2. Roy Hajjar, Ruben AT Mars, and Purna C Kashyap. Harnessing the microbiome for cancer therapy. Nature Reviews Microbiology, pages 1–16, 2026.

3. Sambhawa Priya, Michael B Burns, Tonya Ward, Ruben AT Mars, Beth Adamowicz, Eric F Lock, Purna C Kashyap, Dan Knights, and Ran Blekhman. Identification of shared and disease-specific host gene–microbiome associations across human diseases using multi-omic integration. Nature microbiology, 7(6):780–795, 2022.

4. Maria Kulecka, Jill O’Sullivan, Rachel Fitzgerald, Ana Velikonja, Chloe E Huseyin, Emilio J Laserna-Mendieta, Patricia Ruiz-Limón, Julia Eckenberger, Miriam Vidal-Marín, Bastian-Alexander Truppel, et al. Combining mucosal microbiome and host multi-omics data shows prognostic potential in paediatric ulcerative colitis. Nature Communications, 16(1):7157, 2025.

5. Frank Emmert-Streib, Seppo Parkkila, Reinhard Laubenbacher, Arto Mannermaa, Leroy Hood, and Olli Yli-Harja. The role of digital twins in p4 medicine: A paradigm for modern healthcare. NPJ Digital Medicine, 8(1):735, 2025.

6. Pierfrancesco Novielli, Roberto Bellotti, Mohamad Khalil, Piero Portincasa, and Sabina Tangaro. From networks of data to networks of care in clinical medicine: this is not artificial intelligence. European Journal of Internal Medicine, page 106613, 2025.

7. Lijia Yu, Chunlei Liu, Jean Yee Hwa Yang, and Pengyi Yang. Ensemble deep learning of embeddings for clustering multimodal single-cell omics data. Bioinformatics, 39(6):btad382, 2023.

8. Tal Ashuach, Mariano I Gabitto, Rohan V Koodli, Giuseppe-Antonio Saldi, Michael I Jordan, and Nir Yosef. Multivi: deep generative model for the integration of multimodal data. Nature Methods, 20(8):1222–1231, 2023.

9. Olya Rezaeian, Alparslan Emrah Bayrak, and Onur Asan. Explainability and ai confidence in clinical decision support systems: Effects on trust, diagnostic performance, and cognitive load in breast cancer care. International Journal of Human–Computer Interaction, pages 1–21, 2025.

10. Ricard Argelaguet, Damien Arnol, Danila Bredikhin, Yonatan Deloro, Britta Velten, John C Marioni, and Oliver Stegle. Mofa+: a statistical framework for comprehensive integration of multi-modal single-cell data. Genome biology, 21(1):111, 2020.

11. Qianxing Mo, Sijian Wang, Venkatraman E Seshan, Adam B Olshen, Nikolaus Schultz, Chris Sander, R Scott Powers, Marc Ladanyi, and Ronglai Shen. Pattern discovery and cancer gene identification in integrated cancer genomic data. Proceedings of the National Academy of Sciences, 110(11):4245–4250, 2013.

12. Hamas A Al-Kuhali, Ma Shan, Mohanned Abduljabbar Hael, Eman A Al-Hada, Shamsan A Al-Murisi, Ahmed A Al-Kuhali, Ammar AQ Aldaifl, and Mohammed Elmustafa Amin. Multiview clustering of multi-omics data integration by using a penalty model. BMC bioinformatics, 23(1):288, 2022.

13. Tung Dang, Kie Kumaishi, Erika Usui, Shungo Kobori, Takumi Sato, Yusuke Toda, Yuji Yamasaki, Hisashi Tsujimoto, Yasunori Ichihashi, and Hiroyoshi Iwata. Stochastic variational variable selection for high-dimensional microbiome data. Microbiome, 10(1):236, 2022.

14. Qianxing Mo, Venkatraman E Seshan, Adam B Olshen, Marc Ladanyi, and Ronglai Shen. A fully bayesian latent variable model for integrative clustering of multi-type omics data. Biostatistics, 19(1):71–86, 2018.

15. Eric F Lock and David B Dunson. Bayesian consensus clustering. Bioinformatics, 29(20):2610–2611, 2013.

16. Jalil Taghia, Zhanyu Ma, and Arne Leijon. Bayesian estimation of the von-mises fisher mixture model with variational inference. IEEE transactions on pattern analysis and machine intelligence, 36(9):1701–1715, 2014.

17. Wentao Fan, Wenchuan Zhang, Xiao Dong, and Nizar Bouguila. Clustering-based brain functional segmentation via deep collapsed nonparametric von mises-fisher mixture models. Expert Systems with Applications, page 129739, 2025.

18. Arindam Banerjee, Inderjit S Dhillon, Joydeep Ghosh, Suvrit Sra, and Greg Ridgeway. Clustering on the unit hypersphere using von mises-fisher distributions. Journal of Machine Learning Research, 6(9), 2005.

19. David M Blei, Alp Kucukelbir, and Jon D McAuliffe. Variational inference: A review for statisticians. Journal of the American statistical Association, 112(518):859–877, 2017.

20. Sanket Jantre, Shrijita Bhattacharya, and Tapabrata Maiti. Spike-and-slab shrinkage priors for structurally sparse bayesian neural networks. IEEE Transactions on Neural Networks and Learning Systems, 2024.

21. Tung Dang, Artem Lysenko, Keith A Boroevich, and Tatsuhiko Tsunoda. Vbayesmm: variational bayesian neural network to prioritize important relationships of high-dimensional microbiome multiomics data. Briefings in Bioinformatics, 26(4), 2025.

22. Shinichi Yachida, Sayaka Mizutani, Hirotsugu Shiroma, Satoshi Shiba, Takeshi Nakajima, Taku Sakamoto, Hikaru Watanabe, Keigo Masuda, Yuichiro Nishimoto, Masaru Kubo, et al. Metagenomic and metabolomic analyses reveal distinct stage-specific phenotypes of the gut microbiota in colorectal cancer. Nature medicine, 25(6):968–976, 2019.

23. Anupriya Tripathi, Alexey V Melnik, Jin Xue, Orit Poulsen, Michael J Meehan, Gregory Humphrey, Lingjing Jiang, Gail Ackermann, Daniel McDonald, Dan Zhou, et al. Intermittent hypoxia and hypercapnia, a hallmark of obstructive sleep apnea, alters the gut microbiome and metabolome. Msystems, 3(3):10–1128, 2018.

24. Anders B Dohlman, Diana Arguijo Mendoza, Shengli Ding, Michael Gao, Holly Dressman, Iliyan D Iliev, Steven M Lipkin, and Xiling Shen. The cancer microbiome atlas: a pan-cancer comparative analysis to distinguish tissue-resident microbiota from contaminants. Cell host & microbe, 29(2):281–298, 2021.

25. Lian Narunsky-Haziza, Gregory D Sepich-Poore, Ilana Livyatan, Omer Asraf, Cameron Martino, Deborah Nejman, Nancy Gavert, Jason E Stajich, Guy Amit, Antonio González, et al. Pan-cancer analyses reveal cancer-type-specific fungal ecologies and bacteriome interactions. Cell, 185(20):3789–3806, 2022.

26. Jiacheng Xu and Greg Durrett. Spherical latent spaces for stable variational autoencoders. In Proceedings of the 2018 conference on empirical methods in natural language processing, pages 4503–4513, 2018.

27. Diederik P Kingma. Auto-encoding variational bayes. arXiv preprint arXiv:1312.6114, 2013.

28. Diederik P Kingma and Jimmy Ba. Adam: A method for stochastic optimization. arXiv preprint arXiv:1412.6980, 2014.

29. Wentao Fan and Nizar Bouguila. Online variational learning of generalized dirichlet mixture models with feature selection. Neurocomputing, 126:166–179, 2014.

30. Abraham Gihawi, Yuchen Ge, Jennifer Lu, Daniela Puiu, Amanda Xu, Colin S Cooper, Daniel S Brewer, Mihaela Pertea, and Steven L Salzberg. Major data analysis errors invalidate cancer microbiome findings. MBio, 14(5):e01607–23, 2023.

31. Mengyu He, Ni Zhao, and Glen A Satten. Midasim: a fast and simple simulator for realistic microbiome data. Microbiome, 12(1):135, 2024.

32. Ifeoma Julieth Dikeocha, Abdelkodose Mohammed Al-Kabsi, Hsien-Tai Chiu, and Mohammed Abdullah Alshawsh. Faecalibacterium prausnitzii ameliorates colorectal tumorigenesis and suppresses proliferation of hct116 colorectal cancer cells. Biomedicines, 10(5):1128, 2022.

33. Sarah Obuya, Amr Elkholy, Nagavardhini Avuthu, Michael Behring, Prachi Bajpai, Sumit Agarwal, Hyung-Gyoon Kim, Nefertiti El-Nikhely, Pamela Akinyi, James Orwa, et al. A signature of prevotella copri and faecalibacterium prausnitzii depletion, and a link with bacterial glutamate degradation in the kenyan colorectal cancer patients. Journal of Gastrointestinal Oncology, 13(5):2282, 2022.

34. Satyendra Singh, Ravindra Kumar, Sneha Mittal, Rajesh Sharma, and Ravinder Singh. Butyrate producers, “the sentinel of gut”: Their intestinal significance with and beyond butyrate, and prospective use as microbial therapeutics. Frontiers in Microbiology, 13:1103836, 2023.

35. Hongying Zhang, Qingbin Kong, Jiao Wang, Yangfu Jiang, and Hui Hua. Complex roles of camp–pka–creb signaling in cancer. Experimental hematology & oncology, 9(1):32, 2020.

36. Andong Jin, Fengjing Yang, Hang Li, Geng Wang, Sihua Wang, Song Tong, and Jinbo Gao. 3’-phosphoadenosine 5’-phosphosulfate synthetase 2 (papss2) 1 is a potential diagnostic and prognostic biomarker in colon adenocarcinoma. Scientific Reports, 2026.

37. Sylvia H Duncan, Adela Barcenilla, Colin S Stewart, Susan E Pryde, and Harry J Flint. Acetate utilization 1 and butyryl coenzyme a (coa): acetate-coa transferase in butyrate-producing bacteria from the human large intestine. Applied and environmental microbiology, 68(10):5186–5190, 2002.

38. Alvaro Belenguer, Sylvia H Duncan, A Graham Calder, Grietje Holtrop, Petra Louis, Gerald E Lobley, and Harry J Flint. Two routes of metabolic cross-feeding between bifidobacterium adolescentis and butyrate-producing anaerobes from the human gut. Applied and environmental microbiology, 72(5):3593–3599, 2006.

39. Indhupriya Subramanian, Srikant Verma, Shiva Kumar, Abhay Jere, and Krishanpal Anamika. Multi-omics data integration, interpretation, and its application. Bioinformatics and biology insights, 14:1177932219899051, 2020.

40. Ana R Baião, Zhaoxiang Cai, Rebecca C Poulos, Phillip J Robinson, Roger R Reddel, Qing Zhong, Susana Vinga, and Emanuel Gonçalves. A technical review of multiomics data integration methods: from classical statistical to deep generative approaches. Briefings in bioinformatics, 26(4):bbaf355, 2025.

41. Daniela Mercedes D’Empaire Altimari, Michele Bevere, Elisa Espinet, Yvan Martineau, Elisa Giovannetti, and Víctor Javier Sánchez-Arévalo Lobo. Molecular subtypes in pancreatic cancer: from academic promise to clinical reality. Molecular Cancer, 25(1):134, 2026.

42. Z Gao, D Ghosh, HA Harrington, JG Restrepo, and D Taylor. Dynamics on networks with higher-order interactions. Chaos: An Interdisciplinary Journal of Nonlinear Science, 33(4), 2023.

43. Guilherme Ferraz de Arruda, Alberto Aleta, and Yamir Moreno. Contagion dynamics on higher-order networks. Nature Reviews Physics, 6(8):468–482, 2024.

44. Ke Hu and Linhe Zhu. Pattern dynamics of network information propagation model with higher-order interactions. Information Sciences, page 122812, 2025.

45. Fabian Imdahl, Ehsan Vafadarnejad, Christina Homberger, Antoine-Emmanuel Saliba, and Jörg Vogel. Single-cell rna-sequencing reports growth-condition-specific global transcriptomes of individual bacteria. Nature Microbiology, 5(10):1202–1206, 2020.

46. Bassel Ghaddar, Antara Biswas, Chris Harris, M Bishr Omary, Darren R Carpizo, Martin J Blaser, and Subhajyoti De. Tumor microbiome links cellular programs and immunity in pancreatic cancer. Cancer Cell, 40(10):1240–1253, 2022.

47. Verónica Lloréns-Rico, Joshua A Simcock, Geert RB Huys, and Jeroen Raes. Single-cell approaches in human microbiome research. Cell, 185(15):2725–2738, 2022.

48. Minghui Jia, Senlin Zhu, Ming-Yuan Xue, Hongyi Chen, Jinghong Xu, Mengdi Song, Yifan Tang, Xiaohan Liu, Ye Tao, Tianyu Zhang, et al. Single-cell transcriptomics across 2,534 microbial species reveals functional heterogeneity in the rumen microbiome. Nature Microbiology, 9(7):1884–1898, 2024.

49. Jingjing Li, Yunlong Ma, Yue Cao, Gongwei Zheng, Qing Ren, Cheng Chen, Qunyan Zhu, Yijun Zhou, Yu Lu, Yaru Zhang, et al. Integrating microbial gwas and single-cell transcriptomics reveals associations between host cell populations and the gut microbiome. Nature Microbiology, pages 1–17, 2025.

50. Andrew W Pountain and Itai Yanai. Dissecting microbial communities with single-cell transcriptome analysis. Science, 389(6764):eadp6252, 2025.

