## Supplementary materials for "A hyperspherical deep Bayesian model for interpretable clustering and relationship prediction in microbiome multi-omics integration"

### 1 Supplementary Figures

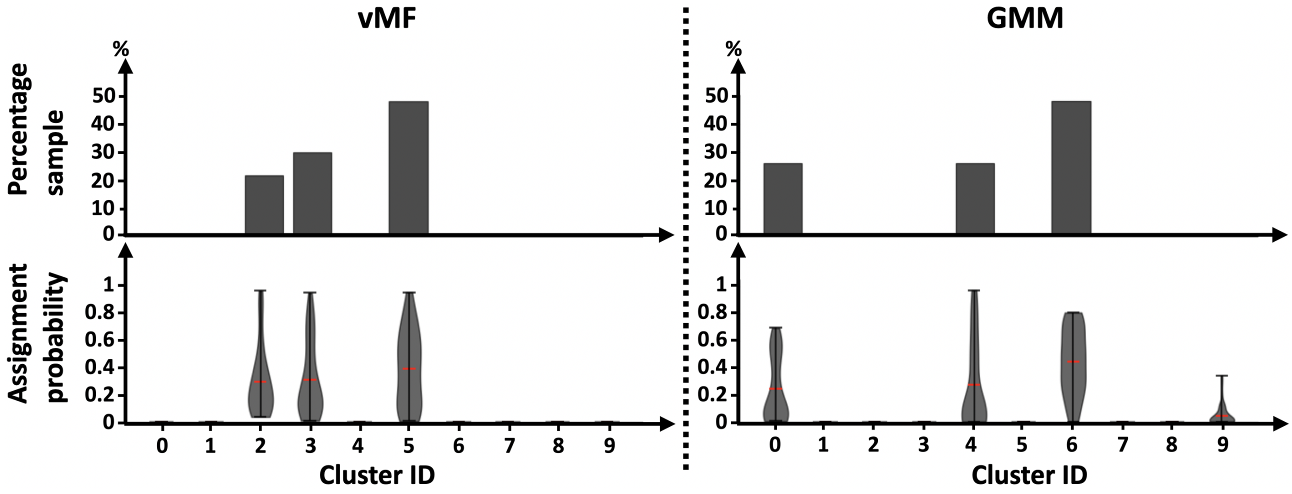


**Supplementary Figure S1**: Probabilistic cluster discovery with uncertainty quantification for three-group structure of dataset A. Top: Percentage of samples assigned to each cluster (0-9). Bottom: Posterior assignment probabilities (violin plots; red lines indicate means) quantifying confidence for each cluster. vMF (von Mises-Fisher mixture model) and GMM (Gaussian mixture model) clustering applied to integrated microbiome-metabolite data.

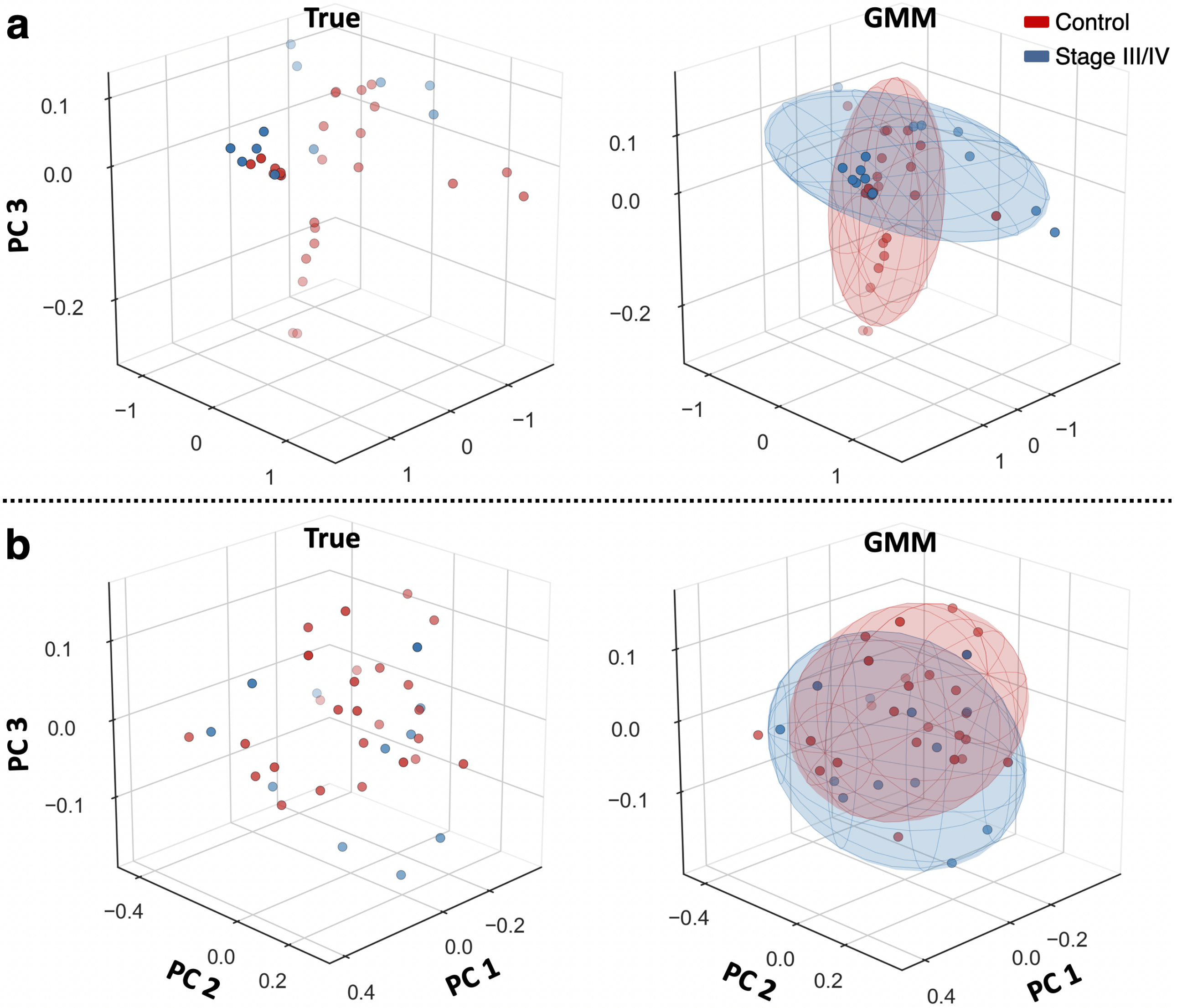


**Supplementary Figure S2**: DBayesCM-GMM latent representation and recovered Gaussian mixture components for the two-group comparison of dataset A. Sample positions are shown as a three-dimensional PCA projection (PC1–PC3) of the Euclidean latent space produced by the standard Gaussian variational autoencoder, with the fitted Gaussian mixture components. "True" panels (left) show the ground-truth partition; "GMM" panels (right) show the recovered components. Red: healthy controls; blue: Stage III/IV cancer. **(a)** microbiome data only. **(b)** microbiome-metabolite data.

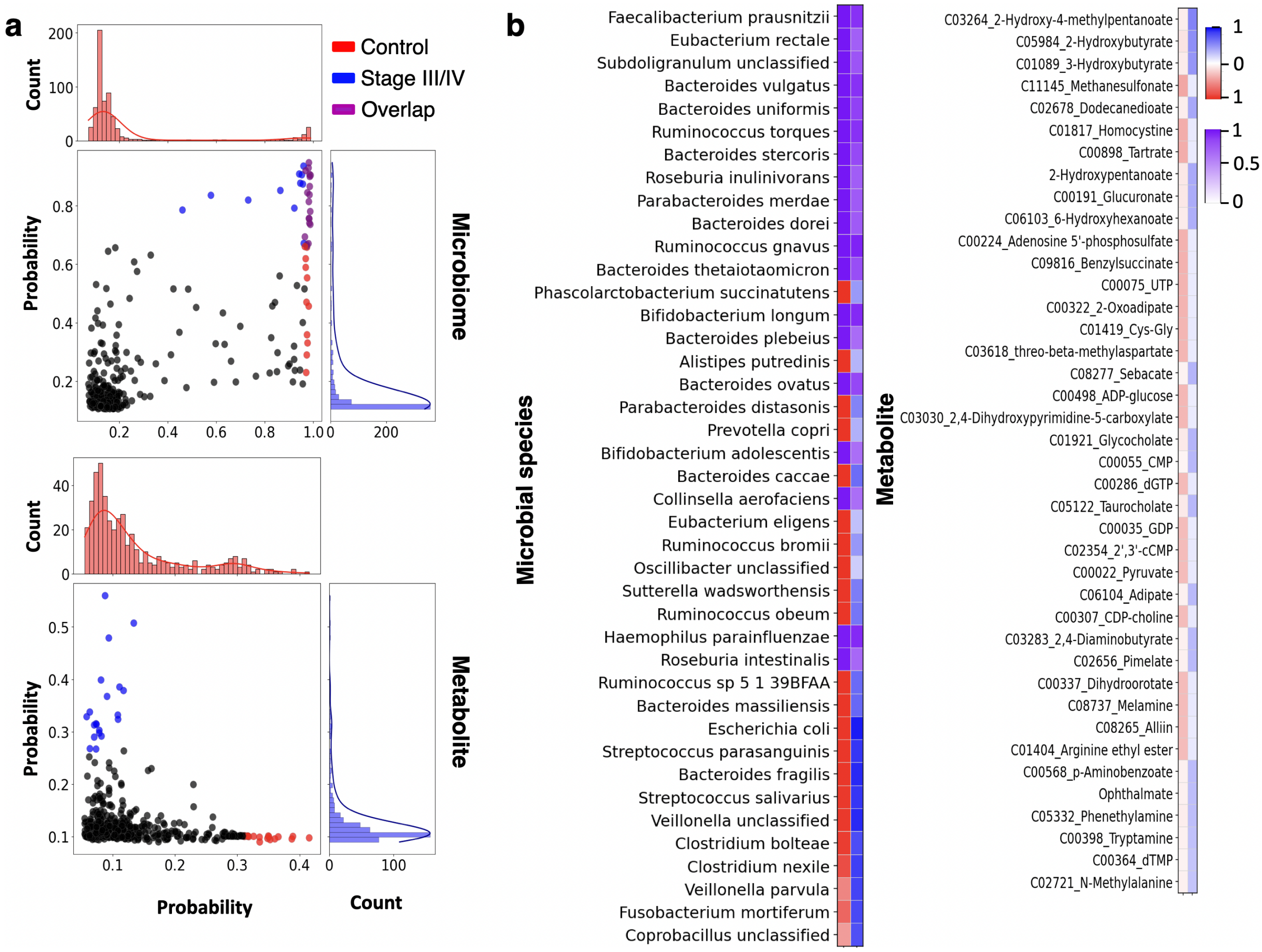


**Supplementary Figure S3**: DBayGenCM achieves superior cancer subtype separation with quantified uncertainty for three-group structure of dataset A. PCA visualization comparing true labels (left) versus clustering predictions from six methods to integrated microbiome-metabolite data. von Mises-Fisher mixture model and Gaussian mixture model display cluster certainty percentages for each subtype (inset boxes). Open circles indicate low-confidence assignments—a unique capability enabling confident clinical decision-making. Red: Stage I/II; Green: Stage III/IV; Blue: Control.

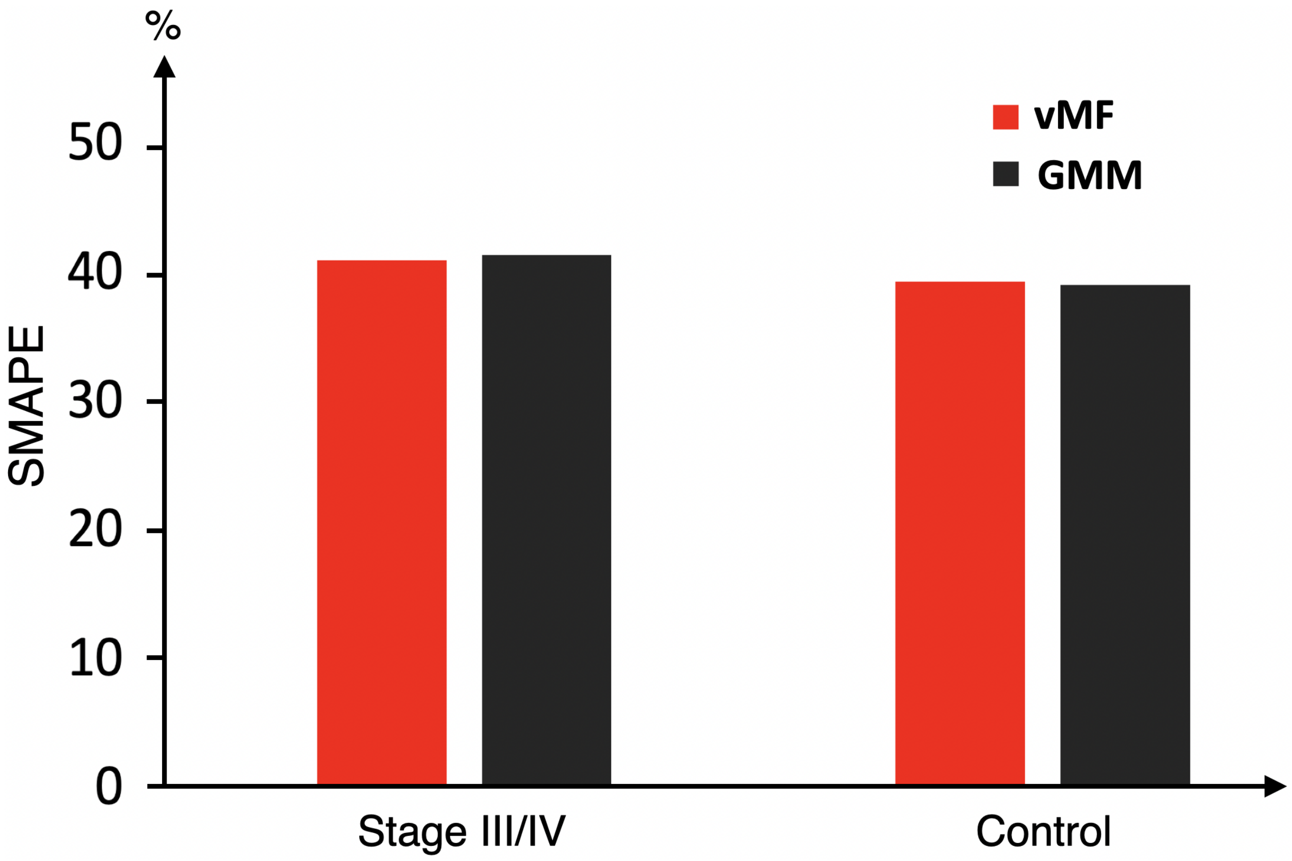


**Supplementary Figure S4**: The SMAPE values are computed for the two approaches of DBayesCM in stage III/IV case and control groups of dataset A.

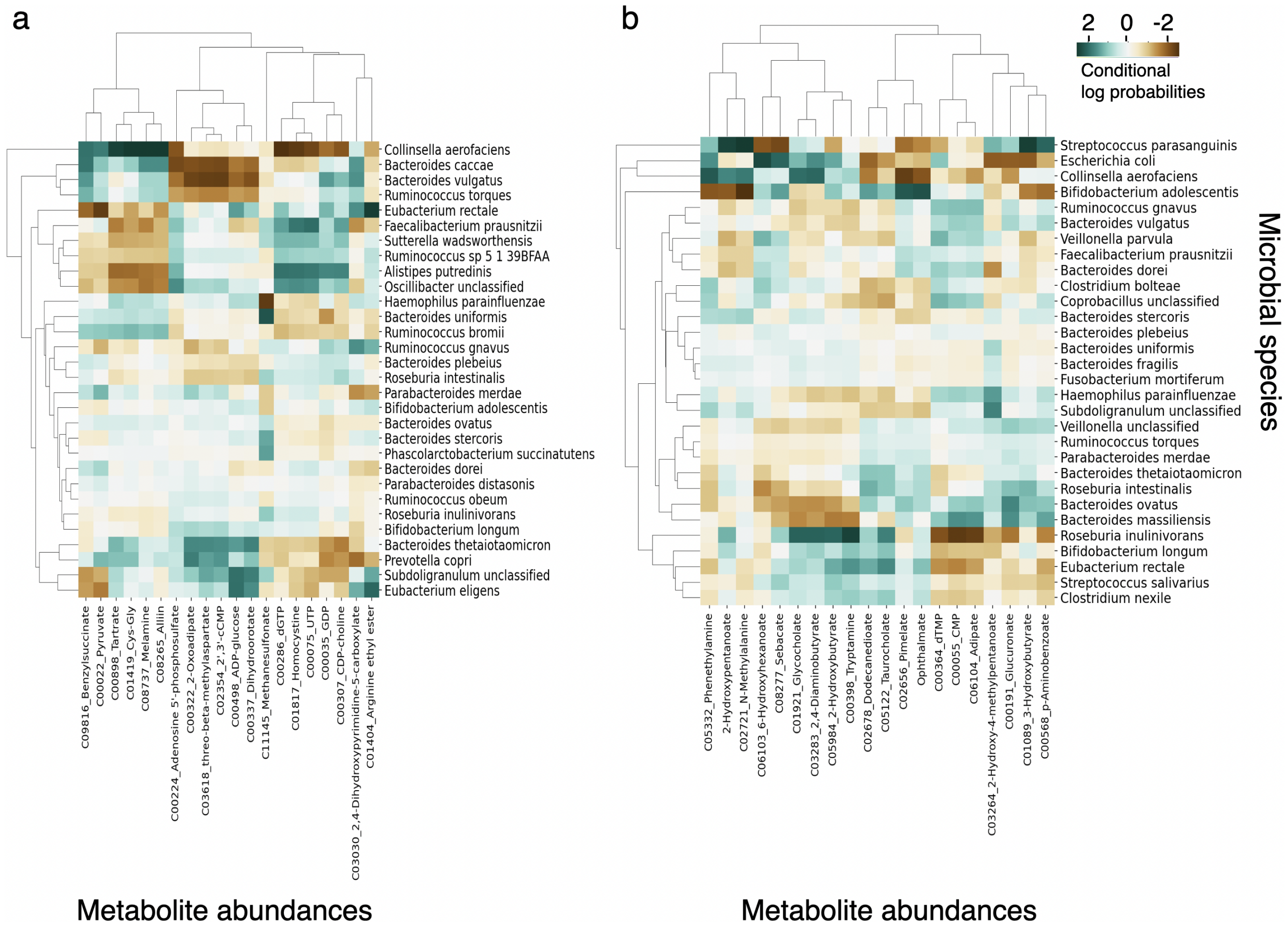


**Supplementary Figure S5**: Heat map of the estimated conditional log probabilities of DBayesCM-GMM for the selected microbial species and metabolite abundances in dataset A. Individual metabolites and microbiomes were hierarchically clustered (Ward's method) using Euclidean distance. (a) stage III/IV. (b) control group.

Table 1: **Supplementary Table S1**:Permutation-null significance of DBayesCM-vMF clustering across all benchmark comparisons. For each comparison, the observed adjusted Rand index (ARI) was tested against a null distribution generated by randomly permuting the true labels ($N = 1000$ permutations). The empirical $p$-value is the fraction of permutations with ARI $\geq$ observed and is reported at the resolution of the test.

| Dataset | Comparison | Observed ARI (mean ± SD) | Null (mean ± SD) | $p$ |
|---|---|---|---|---|
| *Microbiome only* | | | | |
| A (CRC) | Control vs. Stage III/IV | $0.42 \pm 0.040$ | $0.00074 \pm 0.0578$ | $\leq 0.001$ |
| A (CRC) | Control vs. I/II vs. III/IV | $0.30 \pm 0.046$ | $0.00027 \pm 0.0405$ | $\leq 0.001$ |
| B (KIRC) | Stage I vs. III | $0.29 \pm 0.042$ | $0.00021 \pm 0.0245$ | $\leq 0.001$ |
| B (KIRC) | Stage I vs. III vs. IV | $0.15 \pm 0.039$ | $0.00011 \pm 0.0506$ | $\leq 0.001$ |
| C (BRCA) | Stage I vs. IIIA | $0.23 \pm 0.043$ | $0.00012 \pm 0.0237$ | $\leq 0.001$ |
| C (BRCA) | Stage I vs. IIB vs. IIIA | $0.12 \pm 0.034$ | $0.00009 \pm 0.0257$ | $\leq 0.001$ |
| *Microbiome + metabolite (dataset A) / + miRNA (datasets B, C)* | | | | |
| A (CRC) | Control vs. Stage III/IV | $0.51 \pm 0.036$ | $0.00082 \pm 0.0452$ | $\leq 0.001$ |
| A (CRC) | Control vs. I/II vs. III/IV | $0.37 \pm 0.042$ | $0.00035 \pm 0.0550$ | $\leq 0.001$ |
| B (KIRC) | Stage I vs. III | $0.38 \pm 0.038$ | $0.00051 \pm 0.0513$ | $\leq 0.001$ |
| B (KIRC) | Stage I vs. III vs. IV | $0.25 \pm 0.040$ | $0.00019 \pm 0.0263$ | $\leq 0.001$ |
| C (BRCA) | Stage I vs. IIIA | $0.34 \pm 0.034$ | $0.00033 \pm 0.0517$ | $\leq 0.001$ |
| C (BRCA) | Stage I vs. IIB vs. IIIA | $0.22 \pm 0.036$ | $0.00015 \pm 0.0246$ | $\leq 0.001$ |

### 2 Supplementary Methods

#### 2.1 Variational inference for DBayesCM-vMF approach

We expand specifically the Evidence Lower Bound (ELBO) equation as follows:

$$
\begin{aligned}
&\mathcal{L}\left[q\left(\boldsymbol{\Xi}|\Theta\right)\right] = \mathrm{E_q}\left[\log\left(\mathrm{p}\left(\mathbf{X},\boldsymbol{\Xi},\mathbf{Z}\right)\right)\right] - \mathrm{E_q}\left[\log\left(\mathrm{q}\left(\boldsymbol{\Xi},\mathbf{Z}|\Theta\right)\right)\right] \\
&= \mathrm{E_q}\left[\mathrm{logp}\left(\mathbf{X}\right)\right] - \{\mathrm{KL}\left(\mathrm{q}\left(\mathbf{Z}\mid \boldsymbol{Z_\mu},\boldsymbol{Z_\kappa}\right)\|\mathrm{p}\left(\mathbf{Z}\right)\right) + \sum_{\mathrm{k=1}}^{\mathrm{K}}\mathrm{E_q}\left[\log\left(\mathrm{q}\left(\tau_{\mathrm{k}}^{'}|\epsilon_{\mathrm{k}}^{*}\right)\right)\right] - \sum_{\mathrm{k=1}}^{\mathrm{K}}\mathrm{E_q}\left[\log\left(\mathrm{p}\left(\tau_{\mathrm{k}}^{'}|1,\epsilon_{\mathrm{k}}\right)\right)\right] \\
&+ \sum_{n=1}^{N}\sum_{k=1}^{K}\mathrm{E}_q\left[\log\left(q\left(\theta_{nk}|\tau_k^{*}\right)\right)\right] - \sum_{n=1}^{N}\sum_{k=1}^{K}\mathrm{E}_q\left[\log\left(p\left(\theta_{nk}|\tau_k\right)\right)\right] \\
&+ \sum_{n=1}^{N}\sum_{k=1}^{K}\mathrm{E}_q\left[\log\left(q\left(\mu_{nk}|\xi_{nk}^{*},\zeta_{nk}^{*}\kappa_{nk}\right)\right)\right] - \sum_{n=1}^{N}\sum_{k=1}^{K}\mathrm{E}_q\left[\log\left(p\left(\mu_{nk}|\xi_{nk},\zeta_{nk}\kappa_{nk}\right)\right)\right] \\
&+ \sum_{n=1}^{N}\sum_{k=1}^{K}\mathrm{E}_q\left[\log\left(q\left(\kappa_{nk}|\zeta_{nk}^{'*},\zeta_{nk}^{''*}\right)\right)\right] - \sum_{n=1}^{N}\sum_{k=1}^{K}\mathrm{E}_q\left[\log\left(p\left(\kappa_{nk}|\zeta_{nk}^{'},\zeta_{nk}^{''}\right)\right)\right] \\
&+ \sum_{k=1}^{K}\sum_{m=1}^{M}\mathrm{E}_q\left[\log\left(q\left(\psi_{km}|\omega_{km}^{*}\right)\right)\right] - \sum_{k=1}^{K}\sum_{m=1}^{M}\mathrm{E}_q\left[\log\left(p\left(\psi_{km}|\omega_{km}\right)\right)\right] \\
&+ \sum_{k=1}^{K}\sum_{m=1}^{M}\mathrm{E}_q\left[\log\left(q\left(a_{km}|a_{km}^{'*},a_{km}^{''*}\right)\right)\right] - \sum_{k=1}^{K}\sum_{m=1}^{M}\mathrm{E}_q\left[\log\left(p\left(a_{km}|a_{km}^{'},a_{km}^{''}\right)\right)\right] \\
&+ \sum_{k=1}^{K}\sum_{m=1}^{M}\mathrm{E}_q\left[\log\left(q\left(b_{km}|b_{km}^{'*},b_{km}^{''*}\right)\right)\right] - \sum_{k=1}^{K}\sum_{m=1}^{M}\mathrm{E}_q\left[\log\left(p\left(b_{km}|b_{km}^{'},b_{km}^{''}\right)\right)\right] \\
&+ \sum_{k=1}^{K}\sum_{o=1}^{O}\mathrm{E}_q\left[\log\left(q\left(\widetilde{\psi}_{ko}|\widetilde{\omega}_{ko}^{*}\right)\right)\right] - \sum_{k=1}^{K}\sum_{o=1}^{O}\mathrm{E}_q\left[\log\left(p\left(\widetilde{\psi}_{ko}|\widetilde{\omega}_{ko}\right)\right)\right] \\
&+ \sum_{k=1}^{K}\sum_{o=1}^{O}\mathrm{E}_q\left[\log\left(q\left(\widetilde{a}_{ko}|\widetilde{a}_{ko}^{'*},\widetilde{a}_{ko}^{''*}\right)\right)\right] - \sum_{k=1}^{K}\sum_{o=1}^{O}\mathrm{E}_q\left[\log\left(p\left(\widetilde{a}_{ko}|\widetilde{a}_{ko}^{'},\widetilde{a}_{ko}^{''}\right)\right)\right] \\
&+ \sum_{k=1}^{K}\sum_{o=1}^{O}\mathrm{E}_q\left[\log\left(q\left(\widetilde{b}_{ko}|\widetilde{b}_{ko}^{'*},\widetilde{b}_{ko}^{''*}\right)\right)\right] - \sum_{k=1}^{K}\sum_{o=1}^{O}\mathrm{E}_q\left[\log\left(p\left(\widetilde{b}_{ko}|\widetilde{b}_{ko}^{'},\widetilde{b}_{ko}^{''}\right)\right)\right] \\
&+ \mathrm{E}_q\left[\log\left(q\left(e|e^{'*},e^{''*}\right)\right)\right] - \mathrm{E}_q\left[\log\left(p\left(e|e^{'},e^{''}\right)\right)\right] + \mathrm{E}_q\left[\log\left(q\left(f|f^{'*},f^{''*}\right)\right)\right] - \mathrm{E}_q\left[\log\left(p\left(f|f^{'},f^{''}\right)\right)\right] \\
&+ \mathrm{E}_q\left[\log\left(q\left(\widetilde{e}|\widetilde{e}^{'*},\widetilde{e}^{''*}\right)\right)\right] - \mathrm{E}_q\left[\log\left(p\left(\widetilde{e}|\widetilde{e}^{'},\widetilde{e}^{''}\right)\right)\right] + \mathrm{E}_q\left[\log\left(q\left(\widetilde{f}|\widetilde{f}^{'*},\widetilde{f}^{''*}\right)\right)\right] - \mathrm{E}_q\left[\log\left(p\left(\widetilde{f}|\widetilde{f}^{'},\widetilde{f}^{''}\right)\right)\right]\}
\end{aligned} \tag{1}
$$

where $E_q\left[.\right]$ denotes the variational expectations. DBayesCM-vMF implements an infinite Dirichlet process mixture model via truncated stick-breaking construction [1, 2, 3], which automatically determines the optimal number of disease subtypes from data. The stick-breaking process generates mixture weights $\boldsymbol{\tau}$ by sequentially "breaking" a unit-length stick, allocating progressively smaller portions to subsequent clusters until negligible mass remains. The variational posterior over stick-breaking variables factorizes as:

$$q\left(\boldsymbol{\tau}^{'}\right) = \sum_{k=1}^{K} q\left(\tau_k^{'}|\epsilon_k^{*}\right) = \sum_{k=1}^{K} \mathrm{Bernoulli}\left(\sigma(\epsilon_k^{*})\right)), \ \sigma(\epsilon_k^{*})) = \frac{1}{1+e^{-\epsilon_k^{*}}}$$

where $logit_{\epsilon_k^{*}}$ are unconstrained variational parameters learned during training, and $\sigma(.)$ is the sigmoid function. This parameterization enables gradient-based optimization. The KL divergence between posterior and prior for stick-breaking variables is computed via the KL approximation:

$$
\begin{aligned}
\mathrm{KL}\left[q\left(\boldsymbol{\tau}^{'}\right)\|p\left(\boldsymbol{\tau}^{'}\right)\right] &= \sum_{k=1}^{K}\mathrm{E}_q\left[\log\left(q\left(\tau_k^{'}|\epsilon_k^{*}\right)\right)\right] - \sum_{k=1}^{K}\mathrm{E}_q\left[\log\left(p\left(\tau_k^{'}|1,\epsilon_k\right)\right)\right] \\
&= \sum_{k=1}^{K}\left[(1-\tau_k^{'})(\log(1-\tau_k^{'}) - \log(1-\tau_k^{'prior})) + \tau_k^{'}(\log\tau_k^{'} - \log\tau_k^{'prior})\right]
\end{aligned}
$$

where $\tau_k^{'} = \sigma(\epsilon_k^{*})$ is the current posterior mean and $\tau_k^{'prior}$ is sampled from Beta$(1, \epsilon_k)$. This KL term encourages the posterior to match the prior's preference for sparse cluster allocations. The stick-breaking representation generates mixture weights $\tau_k = \tau_k^{'}\prod_{k'=1}^{K}\left(1-\tau_{k'}^{'}\right)$

**Cluster assignment computation $\boldsymbol{\theta}$**
Given joint latent embedding for sample n, cluster responsibilities (soft assignments) are computed as follows:

$$\theta_{nk} = \frac{\exp\left(\mathrm{E}_q\left[\log\tau_k\right] + \mathrm{E}_q\left[\log\left(p\left(Z_n^{Join} \mid \mu_{nk}, \kappa_{nk}\right)\right)\right]\right)}{\sum_{k'=1}^{K}\exp\left(\mathrm{E}_q\left[\log\tau_{k'}\right] + \mathrm{E}_q\left[\log\left(p\left(Z_n^{Join} \mid \mu_{nk'}, \kappa_{nk'}\right)\right)\right]\right)}$$

where:

$$\mathrm{E}_q\left[\log\left(p\left(\boldsymbol{Z}^{Join} \mid \boldsymbol{\mu}, \boldsymbol{\kappa}\right)\right)\right] = \mathrm{E}_q\left[\log\frac{\boldsymbol{\kappa}^{L/2-1}}{(2\pi)^{L/2}\mathcal{I}_{L/2-1}(\boldsymbol{\kappa})}\right] + \mathrm{E}_q[\boldsymbol{\kappa}]\boldsymbol{\mu}^T\boldsymbol{Z}^{Join}$$

Under the Gamma posterior $q\left(\boldsymbol{\kappa} \mid \boldsymbol{\zeta}^{*'}, \boldsymbol{\zeta}^{*''}\right) = \mathcal{G}\left(\boldsymbol{\kappa} \mid \boldsymbol{\zeta}^{*'}, \boldsymbol{\zeta}^{*''}\right)$:

$$\mathrm{E}_q\left[\boldsymbol{\kappa}\right] = \frac{\boldsymbol{\zeta}^{*'}}{\boldsymbol{\zeta}^{*''}},\ \mathrm{E_q}\left[\log\boldsymbol{\kappa}\right] = \psi\left(\boldsymbol{\zeta}^{*'}\right) - \log\boldsymbol{\zeta}^{*''}$$

The KL divergence between posterior and prior for $\boldsymbol{\kappa}$ is computed:

$$\begin{aligned}\mathrm{KL}\left[q\left(\boldsymbol{\kappa}\right)||p\left(\boldsymbol{\kappa}\right)\right] &= \sum_{n=1}^{N}\sum_{k=1}^{K}\mathrm{E}_q\left[\log\left(q\left(\kappa_{nk}|\zeta_{nk}^{'*}, \zeta_{nk}^{''*}\right)\right)\right] - \sum_{n=1}^{N}\sum_{k=1}^{K}\mathrm{E}_q\left[\log\left(p\left(\kappa_{nk}|\zeta_{nk}^{'}, \zeta_{nk}^{''}\right)\right)\right]\\ &= \sum_{n=1}^{N}\sum_{k=1}^{K}\left(\zeta_{nk}^{*'} - \zeta_{nk}^{'}\right)\psi\left(\zeta_{nk}^{*'}\right) - \log\Gamma(\zeta_{nk}^{*'}) + \log\Gamma(\zeta_{nk}^{'}) + \zeta_{nk}^{'}(\log\zeta_{nk}^{*''} - \log\zeta_{nk}^{''}) + \frac{\zeta_{nk}^{*'}(\zeta_{nk}^{''} - \zeta_{nk}^{*''})}{\zeta_{nk}^{*''}}\end{aligned}$$

Under the vMF posterior $q\left(\boldsymbol{\mu} \mid \boldsymbol{\xi}^*, \boldsymbol{\zeta}^*\boldsymbol{\kappa}\right) = \mathcal{V}\left(\boldsymbol{\mu} \mid \boldsymbol{\xi}^*, \boldsymbol{\zeta}^*\boldsymbol{\kappa}\right)$, the expectation over the vMF posterior of $\boldsymbol{\mu}$ is $\mathrm{E}_q\left[\boldsymbol{\mu}\right] = \mathcal{A}_{L/2-1}\left(\boldsymbol{\zeta}^*\mathrm{E}_q(\boldsymbol{\kappa})\right)\boldsymbol{\mu}$. $\mathcal{A}_{L/2-1}\left(\boldsymbol{\kappa}\right) = \mathcal{I}_{L/2}(\boldsymbol{\kappa})/\mathcal{I}_{L/2-1}(\boldsymbol{\kappa})$ is the ratio of Bessel functions, when posterior concentration is large, $\mathcal{A}_{L/2-1}\left(\boldsymbol{\zeta}^*\mathrm{E}_q(\boldsymbol{\kappa})\right) \approx 1$. The KL divergence between posterior and prior for $\boldsymbol{\mu}$ is approximated:

$$\begin{aligned}\mathrm{KL}\left[q\left(\boldsymbol{\mu}\right)||p\left(\boldsymbol{\mu}\right)\right] &= \sum_{n=1}^{N}\sum_{k=1}^{K}\mathrm{E}_q\left[\log\left(q\left(\mu_{nk}|\xi_{nk}^*, \zeta_{nk}^*\kappa_{nk}\right)\right)\right] - \sum_{n=1}^{N}\sum_{k=1}^{K}\mathrm{E}_q\left[\log\left(p\left(\mu_{nk}|\xi_{nk}, \zeta_{nk}\kappa_{nk}\right)\right)\right]\\ &= \sum_{n=1}^{N}\sum_{k=1}^{K}\log\frac{\mathcal{I}_{L/2-1}(\zeta_{nk}\kappa_{nk})}{\mathcal{I}_{L/2-1}(\zeta_{nk}^*\kappa_{nk})} + (L/2-1)(\log\zeta_{nk}^*\kappa_{nk} - \log\zeta_{nk}\kappa_{nk}) - \log\zeta_{nk}^*\kappa_{nk} + \zeta_{nk}\kappa_{nk}\xi_{nk}^{T*}\xi_{nk}\end{aligned}$$

We compute the ELBO for vMF mixtures requires expectations of log-Bessel functions $\mathrm{E}_q\left[\mathcal{I}_{L/2-1}(\boldsymbol{\kappa})\right]$, which lack closed form.

$$\mathrm{E}_q\left[\log\frac{\boldsymbol{\kappa}^{L/2-1}}{(2\pi)^{L/2}\mathcal{I}_{L/2-1}(\boldsymbol{\kappa})}\right] = (L/2-1)\mathrm{E}_q\left[\log\boldsymbol{\kappa}\right] - L/2\log(2\pi) - \mathrm{E}_q\left[\log\left(\mathcal{I}_{L/2-1}(\boldsymbol{\kappa})\right)\right]$$

where $\mathrm{E}_q\left[\log\boldsymbol{\kappa}\right] = \psi\left(\boldsymbol{\zeta}^{*'}\right) - log\boldsymbol{\zeta}^{*''}$. We employ Taylor approximations around expansion points $\widetilde{\boldsymbol{\kappa}}$ as follows[4, 5]:

$$\mathrm{E}_q\left[\log\left(\mathcal{I}_{L/2-1}(\boldsymbol{\kappa})\right)\right] \approx \log\left(\mathcal{I}_{L/2-1}(\widetilde{\boldsymbol{\kappa}})\right) + \frac{\mathcal{I}_{L/2}(\widetilde{\boldsymbol{\kappa}})}{\mathcal{I}_{L/2-1}(\widetilde{\boldsymbol{\kappa}})}\left(\mathrm{E}_q(\boldsymbol{\kappa}) - \widetilde{\boldsymbol{\kappa}}\right)$$

where $\mathrm{E}_q(\boldsymbol{\kappa}) = \frac{\boldsymbol{\zeta}^{*'}}{\boldsymbol{\zeta}^{*''}}$ and $\widetilde{\boldsymbol{\kappa}} = (\boldsymbol{\zeta}^{*'} - 1)/\boldsymbol{\zeta}^{*''}$ if $\boldsymbol{\zeta}^{*'} > 1$, else $\boldsymbol{\zeta}^{*'}/\boldsymbol{\zeta}^{*''}$.
The KL divergence between posterior and prior for $\mathbf{Z}$ is computed [5, 6, 7]:

$$\begin{aligned}\mathrm{KL}\left(q\left(\mathbf{Z} \mid \boldsymbol{Z}_{\boldsymbol{\mu}}, \boldsymbol{Z}_{\boldsymbol{\kappa}}\right)||p\left(\mathbf{Z}\right)\right) &= \boldsymbol{Z}_{\boldsymbol{\mu}}\boldsymbol{Z}_{\boldsymbol{\kappa}}\mathrm{E}_q(\mathbf{Z}) + \log\frac{\boldsymbol{Z}_{\boldsymbol{\kappa}}{}^{L/2-1}}{(2\pi)^{L/2}\mathcal{I}_{L/2-1}(\boldsymbol{Z}_{\boldsymbol{\kappa}})} - \log\left(\frac{2(\pi^{L/2})}{\Gamma(L/2)}\right)^{-1}\\ &= \boldsymbol{Z}_{\boldsymbol{\kappa}}\frac{\mathcal{I}_{L/2}(\boldsymbol{Z}_{\boldsymbol{\kappa}})}{\mathcal{I}_{L/2-1}(\boldsymbol{Z}_{\boldsymbol{\kappa}})} + \left((L/2-1)\log\boldsymbol{Z}_{\boldsymbol{\kappa}} - (L/2)\log(2\pi) - \log\mathcal{I}_{L/2-1}(\boldsymbol{Z}_{\boldsymbol{\kappa}})\right)\\ &+ \frac{L}{2}\log\pi + \log 2 - \log\Gamma(\frac{L}{2})\end{aligned}$$

**Spike-and-slab implementation $\boldsymbol{\gamma}, \boldsymbol{\lambda}$**

For cluster-specific embeddings $\boldsymbol{\gamma}$ (microbiome) and $\boldsymbol{\lambda}$ (omics), spike-and-slab priors enable automatic feature selection through a hierarchical Bayesian structure combining discrete inclusion indicators with continuous shrinkage. For the spike component of the microbiome, the indicator $\psi_{km} \in \{0,1\}$ determines whether taxon m contributes to cluster k. The prior is $p(\psi_{km} \mid \omega_{km}) = \text{Bernoulli}(\omega_{km})$. For differentiability, we use the Gumbel-softmax relaxation [8]:

$$q\left(\psi_{km} \mid \omega_{km}^{*}\right) = \text{Bernoulli}\left(\sigma\left(\text{logit}_{\psi_{km}}\right)\right)$$

where $\text{logit}_{\psi_{km}}$ is a lerned parameter. During forward passes, we sample via:

$$\psi_{km} = \sigma\left(\frac{\text{logit}_{\psi_{km}} + g}{T_g}\right),\ g = \log u - \log(1-u),\ u \sim \text{Uniform}(0,1)$$

where Gumbel temperature $T_g = 0.5$. This provides continuous relaxation enabling backpropagation while approximating discrete sampling. The KL divergence is:

$$\begin{aligned}\text{KL}\left[q\left(\boldsymbol{\psi}\right)||p\left(\boldsymbol{\psi}\right)\right] &= \sum_{k=1}^{K}\sum_{m=1}^{M}\text{E}_q\left[\log\left(q\left(\psi_{km}|\omega_{km}^{*}\right)\right)\right] - \sum_{k=1}^{K}\sum_{m=1}^{M}\text{E}_q\left[\log\left(p\left(\psi_{km}|\omega_{km}\right)\right)\right] \\ &= \sum_{k=1}^{K}\sum_{m=1}^{M}\omega_{km}^{*}\log\frac{\omega_{km}^{*}}{\omega_{km}} + (1-\omega_{km}^{*})\log\frac{1-\omega_{km}^{*}}{1-\omega_{km}}\end{aligned}$$

where $\omega_{km}^{*} = \sigma\left(\text{logit}_{\psi_{km}}\right)$ is the posterior inclusion probability. Local shrinkage parameters $a_{km}, b_{km}$ and global parameters (e,f) control feature-specific and dataset-wide sparsity:

$$\begin{aligned}p(a_{km} \mid a_{km}^{'}, a_{km}^{''}) = \mathcal{G}(1/2,1), &\quad q(a_{km} \mid a_{km}^{'*}, a_{km}^{''*}) = \text{LogNormal}(a_{km}^{'*}, a_{km}^{2''*}) \\ p(b_{km} \mid b_{km}^{'}, b_{km}^{''}) = \mathcal{IG}(1/2,1), &\quad q(b_{km} \mid b_{km}^{'*}, b_{km}^{''*}) = \text{LogNormal}(b_{km}^{'*}, b_{km}^{2''*}) \\ p(e \mid e^{'}, e^{''}) = \mathcal{G}(1/2,1), &\quad q(e \mid e^{'*}, e^{''*}) = \text{LogNormal}(\mid e^{'*}, e^{2''*}) \\ p(f \mid f^{'}, f^{''}) = \mathcal{IG}(1/2,1), &\quad q(f \mid f^{'*}, f^{''*}) = \text{LogNormal}(f^{'*}, f^{2''*})\end{aligned}$$

The KL divergence between posterior and prior for $\mathbf{a}$ is computed:

$$\begin{aligned}\text{KL}[q(\mathbf{a})|p(\mathbf{a})] &= \sum_{k=1}^{K}\sum_{m=1}^{M}\text{E}_q\left[\log\left(q\left(a_{km}|a_{km}^{'*}, a_{km}^{''*}\right)\right)\right] - \sum_{k=1}^{K}\sum_{m=1}^{M}\text{E}_q\left[\log\left(p\left(a_{km}|a_{km}^{'}, a_{km}^{''}\right)\right)\right] \\ &\approx \sum_{k=1}^{K}\sum_{m=1}^{M}(a_{km}^{'}-1)\text{E}_q[\log a_{km}] - a_{km}^{''}\text{E}_q[a_{km}] + a_{km}^{'}\log a_{km}^{''} - \log\Gamma(a_{km}^{'}) + \frac{1}{2}\log(2\pi a_{km}^{2''*}) + \frac{1}{2} \\ &\approx \sum_{k=1}^{K}\sum_{m=1}^{M}(a_{km}^{'}-1)a_{km}^{'*} - a_{km}^{''}e^{a_{km}^{'*}+a_{km}^{2''*}/2} + a_{km}^{'}\log a_{km}^{''} - \log\Gamma(a_{km}^{'}) + \frac{1}{2}\log(2\pi a_{km}^{2''*}) + \frac{1}{2}\end{aligned}$$

where $\text{E}_q[\log a_{km}] = a_{km}^{'*}$ and $\text{E}_q[a_{km}] = e^{a_{km}^{'*}+a_{km}^{2''*}/2}$.

The KL divergence between posterior and prior for $\mathbf{b}$ is computed:

$$\begin{aligned}\text{KL}[q(\mathbf{b})|p(\mathbf{b})] &= \sum_{k=1}^{K}\sum_{m=1}^{M}\text{E}_q\left[\log\left(q\left(b_{km}|b_{km}^{'*}, b_{km}^{''*}\right)\right)\right] - \sum_{k=1}^{K}\sum_{m=1}^{M}\text{E}_q\left[\log\left(p\left(b_{km}|b_{km}^{'}, b_{km}^{''}\right)\right)\right] \\ &\approx \sum_{k=1}^{K}\sum_{m=1}^{M}-(b_{km}^{'}+1)b_{km}^{'*} - b_{km}^{''}e^{-b_{km}^{'*}+b_{km}^{2''*}/2} - b_{km}^{'}\log b_{km}^{''} + \log\Gamma(b_{km}^{'}) + \frac{1}{2}\log(2\pi b_{km}^{2''*}) + \frac{1}{2}\end{aligned}$$

The KL divergence between posterior and prior for $\mathbf{e}$ is computed:

$$\begin{aligned}\text{KL}[q(\mathbf{e})|p(\mathbf{e})] &= \text{E}_q\left[\log\left(q\left(e|e^{'*}, e^{''*}\right)\right)\right] - \text{E}_q\left[\log\left(p\left(e|e^{'}, e^{''}\right)\right)\right] \\ &\approx (e^{'}-1)e^{'*} - e^{''}e^{e_{km}^{'*}+e^{2''*}/2} + e^{'}\log e^{''} - \log\Gamma(e^{'}) + \frac{1}{2}\log(2\pi e^{2''*}) + \frac{1}{2}\end{aligned}$$

The KL divergence between posterior and prior for $\mathbf{f}$ is computed:

$$\begin{aligned}\mathrm{KL}[q(\mathbf{f})|p(\mathbf{f})] &= \mathrm{E}_q\left[\log\left(q\left(f|f^{'*}, f^{''*}\right)\right)\right] - \mathrm{E}_q\left[\log\left(p\left(f|f^{'}, f^{''}\right)\right)\right] \\ &\approx -(f^{'}+1)f^{'*} - f^{''}e^{-f^{'*}+f^{2''*}/2} - f^{'}\log f^{''} + \log\Gamma(f^{'}) + \frac{1}{2}\log(2\pi f^{2''*}) + \frac{1}{2}\end{aligned}$$

For the host omics embedding $\boldsymbol{\lambda}$, we employed a similar approach for all hyperparameters. The KL divergences are computed as follows:

$$\begin{aligned}\mathrm{KL}\left[q\left(\widetilde{\boldsymbol{\psi}}\right)||p\left(\widetilde{\boldsymbol{\psi}}\right)\right] &= \sum_{k=1}^{K}\sum_{m=1}^{M}\mathrm{E}_q\left[\log\left(q\left(\widetilde{\psi_{km}}|\widetilde{\omega_{km}}^{*}\right)\right)\right] - \sum_{k=1}^{K}\sum_{m=1}^{M}\mathrm{E}_q\left[\log\left(p\left(\widetilde{\psi_{km}}|\widetilde{\omega_{km}}\right)\right)\right] \\ &= \sum_{k=1}^{K}\sum_{m=1}^{M}\widetilde{\omega_{km}}^{*}\log\frac{\widetilde{\omega_{km}}^{*}}{\widetilde{\omega_{km}}} + (1-\widetilde{\omega_{km}}^{*})\log\frac{1-\widetilde{\omega_{km}}^{*}}{1-\widetilde{\omega_{km}}}\end{aligned}$$

$$\begin{aligned}\mathrm{KL}[q(\widetilde{\mathbf{a}})|p(\widetilde{\mathbf{a}})] &= \sum_{k=1}^{K}\sum_{m=1}^{M}\mathrm{E}_q\left[\log\left(q\left(\widetilde{a}_{km}|\widetilde{a_{km}}^{'*}, \widetilde{a_{km}}^{''*}\right)\right)\right] - \sum_{k=1}^{K}\sum_{m=1}^{M}\mathrm{E}_q\left[\log\left(p\left(\widetilde{a_{km}}|\widetilde{a_{km}}^{'}, \widetilde{a_{km}}^{''}\right)\right)\right] \\ &\approx \sum_{k=1}^{K}\sum_{m=1}^{M}(\widetilde{a_{km}}^{'}-1)\widetilde{a^{'*}_{km}} - \widetilde{a^{''}_{km}}e^{\widetilde{a^{'*}_{km}}+\widetilde{a^{2''*}_{km}}/2} + \widetilde{a^{'}_{km}}\log\widetilde{a^{''}_{km}} - \log\Gamma(\widetilde{a_{km}}^{'}) + \frac{1}{2}\log(2\pi\widetilde{a^{2''*}_{km}}) + \frac{1}{2}\end{aligned}$$

$$\begin{aligned}\mathrm{KL}[q(\widetilde{\mathbf{b}})|p(\widetilde{\mathbf{b}})] &= \sum_{k=1}^{K}\sum_{m=1}^{M}\mathrm{E}_q\left[\log\left(q\left(\widetilde{b_{km}}|\widetilde{b_{km}}^{'*}, \widetilde{b_{km}}^{''*}\right)\right)\right] - \sum_{k=1}^{K}\sum_{m=1}^{M}\mathrm{E}_q\left[\log\left(p\left(\widetilde{b_{km}}|\widetilde{b_{km}}^{'}, \widetilde{b_{km}}^{''}\right)\right)\right] \\ &\approx \sum_{k=1}^{K}\sum_{m=1}^{M} -(\widetilde{b_{km}}^{'}+1)\widetilde{b_{km}}^{'*} - \widetilde{b_{km}}^{''}e^{-\widetilde{b_{km}}^{'*}+\widetilde{b_{km}}^{2''*}/2} - \widetilde{b_{km}}^{'}\log\widetilde{b_{km}}^{''} + \log\Gamma(\widetilde{b_{km}}^{'}) + \frac{1}{2}\log(2\pi\widetilde{b_{km}}^{2''*}) + \frac{1}{2}\end{aligned}$$

$$\begin{aligned}\mathrm{KL}[q(\widetilde{\mathbf{e}})|p(\widetilde{\mathbf{e}})] &= \mathrm{E}_q\left[\log\left(q\left(\widetilde{e}|\widetilde{e}^{'*}, \widetilde{e}^{''*}\right)\right)\right] - \mathrm{E}_q\left[\log\left(p\left(\widetilde{e}|\widetilde{e}^{'}, \widetilde{e}^{''}\right)\right)\right] \\ &\approx (\widetilde{e}^{'}-1)\widetilde{e}^{'*} - \widetilde{e}^{''}e^{\widetilde{e}^{'*}_{km}+\widetilde{e}^{2''*}/2} + \widetilde{e}^{'}\log\widetilde{e}^{''} - \log\Gamma(\widetilde{e}^{'}) + \frac{1}{2}\log(2\pi\widetilde{e}^{2''*}) + \frac{1}{2}\end{aligned}$$

$$\begin{aligned}\mathrm{KL}[q(\widetilde{\mathbf{f}})|p(\widetilde{\mathbf{f}})] &= \mathrm{E}_q\left[\log\left(q\left(\widetilde{f}|\widetilde{f}^{'*}, \widetilde{f}^{''*}\right)\right)\right] - \mathrm{E}_q\left[\log\left(p\left(\widetilde{f}|\widetilde{f}^{'}, \widetilde{f}^{''}\right)\right)\right] \\ &\approx -(\widetilde{f}^{'}+1)\widetilde{f}^{'*} - \widetilde{f}^{''}e^{-\widetilde{f}^{'*}+\widetilde{f}^{2''*}/2} - \widetilde{f}^{'}\log\widetilde{f}^{''} + \log\Gamma(\widetilde{f}^{'}) + \frac{1}{2}\log(2\pi\widetilde{f}^{2''*}) + \frac{1}{2}\end{aligned}$$

#### 2.2 Variational inference for DBayesCM-GMM approach

We expand specifically the Evidence Lower Bound (ELBO) equation as follows:

$$
\begin{aligned}
\mathcal{L}\left[q\left(\boldsymbol{\Xi}|\Theta\right)\right] &= \mathrm{E_q}\left[\log\left(\mathrm{p}\left(\mathbf{X},\boldsymbol{\Xi},\mathbf{Z}\right)\right)\right] - \mathrm{E_q}\left[\log\left(\mathrm{q}\left(\boldsymbol{\Xi},\mathbf{Z}|\Theta\right)\right)\right] \\
&= \mathrm{E_q}\left[\mathrm{logp}\left(\mathbf{X}\right)\right] - \{\mathrm{KL}\left(\mathrm{q}\left(\mathbf{Z}\mid \boldsymbol{Z_\mu},\boldsymbol{Z_\kappa}\right)\|\mathrm{p}\left(\mathbf{Z}\right)\right) + \sum_{\mathrm{k=1}}^{\mathrm{K}} \mathrm{E_q}\left[\log\left(\mathrm{q}\left(\eta_{\mathrm{k}}^{'}|\varepsilon_{\mathrm{k}}^{*}\right)\right)\right] - \sum_{\mathrm{k=1}}^{\mathrm{K}} \mathrm{E_q}\left[\log\left(\mathrm{p}\left(\eta_{\mathrm{k}}^{'}|1,\varepsilon_{\mathrm{k}}\right)\right)\right] \\
&+ \sum_{n=1}^{N}\sum_{k=1}^{K} \mathrm{E}_q\left[\log\left(q\left(\theta_{nk}|\eta_k^*\right)\right)\right] - \sum_{n=1}^{N}\sum_{k=1}^{K} \mathrm{E}_q\left[\log\left(p\left(\theta_{nk}|\eta_k\right)\right)\right] \\
&+ \sum_{n=1}^{N}\sum_{k=1}^{K} \mathrm{E}_q\left[\log\left(q\left(\rho_{nk}|\varrho_{nk}^{*},\vartheta_{nk}^{2*}\right)\right)\right] - \sum_{n=1}^{N}\sum_{k=1}^{K} \mathrm{E}_q\left[\log\left(p\left(\rho_{nk}|\varrho_{nk},\vartheta_{nk}^{2}\right)\right)\right] \\
&+ \sum_{n=1}^{N}\sum_{k=1}^{K} \mathrm{E}_q\left[\log\left(q\left(\sigma_{nk}^{2}|c_{nk}^{*},d_{nk}^{*}\right)\right)\right] - \sum_{n=1}^{N}\sum_{k=1}^{K} \mathrm{E}_q\left[\log\left(p\left(\sigma_{nk}^{2}|c_{nk},d_{nk}\right)\right)\right] \\
&+ \sum_{k=1}^{K}\sum_{m=1}^{M} \mathrm{E}_q\left[\log\left(q\left(\psi_{km}|\omega_{km}^{*}\right)\right)\right] - \sum_{k=1}^{K}\sum_{m=1}^{M} \mathrm{E}_q\left[\log\left(p\left(\psi_{km}|\omega_{km}\right)\right)\right] \\
&+ \sum_{k=1}^{K}\sum_{m=1}^{M} \mathrm{E}_q\left[\log\left(q\left(a_{km}|a_{km}^{'*},a_{km}^{''*}\right)\right)\right] - \sum_{k=1}^{K}\sum_{m=1}^{M} \mathrm{E}_q\left[\log\left(p\left(a_{km}|a_{km}^{'},a_{km}^{''}\right)\right)\right] \\
&+ \sum_{k=1}^{K}\sum_{m=1}^{M} \mathrm{E}_q\left[\log\left(q\left(b_{km}|b_{km}^{'*},b_{km}^{''*}\right)\right)\right] - \sum_{k=1}^{K}\sum_{m=1}^{M} \mathrm{E}_q\left[\log\left(p\left(b_{km}|b_{km}^{'},b_{km}^{''}\right)\right)\right] \\
&+ \sum_{k=1}^{K}\sum_{o=1}^{O} \mathrm{E}_q\left[\log\left(q\left(\widetilde{\psi}_{ko}|\widetilde{\omega}_{ko}^{*}\right)\right)\right] - \sum_{k=1}^{K}\sum_{o=1}^{O} \mathrm{E}_q\left[\log\left(p\left(\widetilde{\psi}_{ko}|\widetilde{\omega}_{ko}\right)\right)\right] \\
&+ \sum_{k=1}^{K}\sum_{o=1}^{O} \mathrm{E}_q\left[\log\left(q\left(\widetilde{a}_{ko}|\widetilde{a}_{ko}^{'*},\widetilde{a}_{ko}^{''*}\right)\right)\right] - \sum_{k=1}^{K}\sum_{o=1}^{O} \mathrm{E}_q\left[\log\left(p\left(\widetilde{a}_{ko}|\widetilde{a}_{ko}^{'},\widetilde{a}_{ko}^{''}\right)\right)\right] \\
&+ \sum_{k=1}^{K}\sum_{o=1}^{O} \mathrm{E}_q\left[\log\left(q\left(\widetilde{b}_{ko}|\widetilde{b}_{ko}^{'*},\widetilde{b}_{ko}^{''*}\right)\right)\right] - \sum_{k=1}^{K}\sum_{o=1}^{O} \mathrm{E}_q\left[\log\left(p\left(\widetilde{b}_{ko}|\widetilde{b}_{ko}^{'},\widetilde{b}_{ko}^{''}\right)\right)\right] \\
&+ \mathrm{E}_q\left[\log\left(q\left(e|e^{'*},e^{''*}\right)\right)\right] - \mathrm{E}_q\left[\log\left(p\left(e|e^{'},e^{''}\right)\right)\right] + \mathrm{E}_q\left[\log\left(q\left(f|f^{'*},f^{''*}\right)\right)\right] - \mathrm{E}_q\left[\log\left(p\left(f|f^{'},f^{''}\right)\right)\right] \\
&+ \mathrm{E}_q\left[\log\left(q\left(\widetilde{e}|\widetilde{e}^{'*},\widetilde{e}^{''*}\right)\right)\right] - \mathrm{E}_q\left[\log\left(p\left(\widetilde{e}|\widetilde{e}^{'},\widetilde{e}^{''}\right)\right)\right] + \mathrm{E}_q\left[\log\left(q\left(\widetilde{f}|\widetilde{f}^{'*},\widetilde{f}^{''*}\right)\right)\right] - \mathrm{E}_q\left[\log\left(p\left(\widetilde{f}|\widetilde{f}^{'},\widetilde{f}^{''}\right)\right)\right]\}
\end{aligned}
\tag{2}
$$

In this context, $E_q[.]$ represents the variational expectations. The DBayesCM-GMM employs an infinite Dirichlet process mixture model through a truncated stick-breaking construction, utilizing Gaussian mixture posteriors to define cluster geometries. The spike-and-slab methodology for sparse feature selection is analogous to that used in DBayesCM-vMF.

Given joint latent embedding for sample n, cluster responsibilities (soft assignments) are computed as follows [1, 2]:

$$
\theta_{nk} = \frac{\exp\left(\mathrm{E}_q\left[\log\eta_k\right] + \mathrm{E}_q\left[\log\left(\mathcal{N}\left(Z_n^{Join} \mid \rho_{nk},\sigma_{nk}^2\right)\right)\right]\right)}{\sum\limits_{k'=1}^{K} \exp\left(\mathrm{E}_q\left[\log\eta_{k'}\right] + \mathrm{E}_q\left[\log\left(\mathcal{N}\left(Z_n^{Join} \mid \rho_{nk'},\sigma_{nk'}^2\right)\right)\right]\right)}
$$

The KL divergence between the variational posterior and the stick-breaking prior is computed as follows:

$$
\begin{aligned}
\mathrm{KL}\left[q\left(\boldsymbol{\eta}^{'}\right)\|p\left(\boldsymbol{\eta}^{'}\right)\right] &= \sum_{k=1}^{K} \mathrm{E}_q\left[\log\left(q\left(\eta_k^{'}|\varepsilon_k^{*}\right)\right)\right] - \sum_{k=1}^{K} \mathrm{E}_q\left[\log\left(p\left(\eta_k^{'}|1,\varepsilon_k\right)\right)\right] \\
&= \sum_{k=1}^{K}\left[(1-\eta_k^{'})(\log(1-\eta_k^{'}) - \log(1-\eta_k^{'prior})) + \eta_k^{'}(\log\eta_k^{'} - \log\eta_k^{'prior})\right]
\end{aligned}
$$

In this context, $\eta_k^{'} = \sigma(\varepsilon_k^{*})$ represents the current posterior mean, while $\eta_k^{'prior}$ is drawn from a Beta$(1,\varepsilon_k)$ distribution. This KL divergence term serves to align the posterior distribution with the prior's inclination towards sparse cluster allocations. Under the Normal posterior $q\left(\rho_{nk}|\varrho_{nk}^{*},\vartheta_{nk}^{2*}\right) = \mathcal{N}\left(\rho_{nk}|\varrho_{nk}^{*},\vartheta_{nk}^{2*}\right)$, the KL divergence between posterior and prior for $\boldsymbol{\rho}$ is computed:

$$\mathrm{KL}[q(\boldsymbol{\rho})|p(\boldsymbol{\rho})] = \sum_{n=1}^{N}\sum_{k=1}^{K}\mathrm{E}_q\left[\log\left(q\left(\rho_{nk}|\varrho^*_{nk},\vartheta^{2*}_{nk}\right)\right)\right] - \sum_{n=1}^{N}\sum_{k=1}^{K}\mathrm{E}_q\left[\log\left(p\left(\rho_{nk}|\varrho_{nk},\vartheta^{2}_{nk}\right)\right)\right]$$
$$= \sum_{n=1}^{N}\sum_{k=1}^{K}\frac{1}{2}\left[\log\frac{\vartheta^2_{nk}}{\vartheta^{2*}_{nk}} + \frac{\vartheta^{2*}_{nk} + (\varrho^*_{nk} - \varrho_{nk})^2}{\vartheta^2_{nk}} - 1\right]$$

Under the Inverse-Gamma posterior $q\left(\sigma^2_{nk}|c^*_{nk},d^*_{nk}\right) = \mathcal{IG}\left(\sigma^2_{nk}|c^*_{nk},d^*_{nk}\right)$, the KL divergence between posterior and prior for $\boldsymbol{\sigma^2}$ is computed:

$$\mathrm{KL}[q(\boldsymbol{\sigma^2})|p(\boldsymbol{\sigma^2})] = \sum_{n=1}^{N}\sum_{k=1}^{K}\mathrm{E}_q\left[\log\left(q\left(\sigma^2_{nk}|c^*_{nk},d^*_{nk}\right)\right)\right] - \sum_{n=1}^{N}\sum_{k=1}^{K}\mathrm{E}_q\left[\log\left(p\left(\sigma^2_{nk}|c_{nk},d_{nk}\right)\right)\right]$$
$$= c_{nk}\log\frac{d^*_{nk}}{d_{nk}} + (c^*_{nk} - c_{nk})\psi(\alpha_{c^*_{nk}}) - \log\Gamma(c^*_{nk}) + \log\Gamma(c_{nk}) + c^*_{nk}\left(\frac{d_{nk}}{d^*_{nk}} - 1\right)$$

where $\psi(.) = \frac{d\log\Gamma(.)}{dx}$ represents the digamma function. This KL divergence term imposes a penalty on deviations from the unit-scale prior, thereby promoting automatic shrinkage of cluster variances unless the data necessitate broader distributions.

### References


[1] D. M. Blei and M. I. Jordan, "Variational inference for dirichlet process mixtures," 2006.

[2] D. M. Blei, A. Kucukelbir, and J. D. McAuliffe, "Variational inference: A review for statisticians," *Journal of the American statistical Association*, vol. 112, no. 518, pp. 859–877, 2017.

[3] T. Dang, K. Kumaishi, E. Usui, S. Kobori, T. Sato, Y. Toda, Y. Yamasaki, H. Tsujimoto, Y. Ichihashi, and H. Iwata, "Stochastic variational variable selection for high-dimensional microbiome data," *Microbiome*, vol. 10, no. 1, p. 236, 2022.

[4] J. Taghia, Z. Ma, and A. Leijon, "Bayesian estimation of the von-mises fisher mixture model with variational inference," *IEEE transactions on pattern analysis and machine intelligence*, vol. 36, no. 9, pp. 1701–1715, 2014.

[5] W. Fan, W. Zhang, X. Dong, and N. Bouguila, "Clustering-based brain functional segmentation via deep collapsed nonparametric von mises-fisher mixture models," *Expert Systems with Applications*, p. 129739, 2025.

[6] J. Xu and G. Durrett, "Spherical latent spaces for stable variational autoencoders," in *Proceedings of the 2018 conference on empirical methods in natural language processing*, 2018, pp. 4503–4513.

[7] T. R. Davidson, L. Falorsi, N. De Cao, T. Kipf, and J. M. Tomczak, "Hyperspherical variational auto-encoders," *arXiv preprint arXiv:1804.00891*, 2018.

[8] S. Jantre, S. Bhattacharya, and T. Maiti, "Spike-and-slab shrinkage priors for structurally sparse bayesian neural networks," *IEEE Transactions on Neural Networks and Learning Systems*, vol. 36, no. 6, pp. 11 176–11 188, 2024.
